# Social isolation upregulates *takeout* expression in female *Drosophila melanogaster* to promote sucrose feeding

**DOI:** 10.64898/2025.12.09.693125

**Authors:** Elisabeth A. Tawa, Lisa Schneper, Katherine I. Cho, Mindy Sim, Eve Schoeffler, Daniel A. Notterman

## Abstract

*Drosophila melanogaster* provides a model system to examine how environmental stress interacts with sex to induce changes in brain function and behavior. Previous research suggests that social isolation induces changes in gene expression that encode a starvation-like brain state and reduce sleep. However, the extent to which social isolation alters behaviors via sex-specific brain changes is unclear. Here, we use *Drosophila melanogaster* to explore sex differences in chronic social isolation-induced behavioral and transcriptomic changes. We focused on *takeout* (*to*), a gene encoding a putative juvenile hormone-binding protein, as a target that is upregulated solely in females following social isolation. Male and female adult flies were exposed to chronic social isolation, and multiple behavioral sex differences were identified through tests of activity, motivation, aggression, and sugar consumption. RNA-seq analysis also identified several candidate genes that were associated with sex differences in isolation-induced behavioral changes. Our findings suggest that social isolation is sufficiently stressful to reveal latent sex differences in behavior, despite having no impact on survival. *To* expression and sucrose consumption were upregulated exclusively in females following social isolation. Following *to* knockdown in *to*-expressing cells, sucrose consumption decreased in socially isolated females but increased in males. However, *to* knock down, *to* overexpression, and *transformer* knock down in neurons did not change sucrose-feeding behavior between control and isolated females. Overall, our results suggest that manipulating *to* expression influences sucrose-feeding in opposite directions between females and males following social isolation, and that isolation-induced *to* overexpression in non-neuronal cells in the brain or head may play a role in communicating information about females’ nutritional status to the brain. Additional roles for *to* in stress-related behaviors and behavioral sex differences should be explored, as well as whether *to* participates in signaling pathways that may be functionally conserved in human disorders.

**Author Summary:** Chronic stress contributes to detrimental health effects, but our understanding of how stress induces sex differences in brain gene expression and behavior is incomplete. Here, we use a combination of behavioral testing, RNA sequencing, and genetic manipulations in *Drosophila melanogaster* to explore how social isolation reveals latent sex differences in gene expression and stress-relevant behaviors. We found the most pronounced sex differences in behaviors related to feeding and motivation. RNA profiling revealed isolated female-specific upregulation of over 100 genes, with many of them relating to reproduction and energy metabolism. We manipulated expression of the candidate gene *takeout (to)* and found that downregulating *to* in all *to*-expressing cells decreases sucrose-feeding in isolated females but increases it in isolated males. Our results suggest that within a chronic stress context, sex-specific effectors in the head may regulate gene expression related to feeding and macronutrient choice to ensure that females prioritize survival over reproduction. Learning more about this system in flies could provide insight into functionally analogous pathways in humans that may be dysregulated in female-biased stress-related disorders.

## Introduction

Social isolation has emerged in recent decades as a worldwide public health issue. The World Health Organization estimates that one in four older adults and five to fifteen percent of adolescents experience social isolation and loneliness globally (1) while the Centers for Disease Control and Prevention estimates that one in three adults experience these problems in the United States (2). In humans, isolation increases the risk of early death and is associated with several diseases and psychiatric conditions, including cardiovascular disease, stroke, type 2 diabetes, depression, anxiety, suicidality, and dementia (1,2). Social isolation has also been found to enhance aggression, depression, and anxiety in humans undergoing solitary confinement (3,4).

Common social stressors across animal species include maternal separation, social deprivation, reduced parental care, and aggression from caregivers early in life (5). In adulthood, social stressors manifest as competition for mates and territory, within-group competition for resources and social rank, and between-group conflict (5). Individuals within a social group may benefit the most in terms of fitness by living in a group of intermediate size. This provides safety in numbers and allows for engagement in cooperative behaviors. If the social group is too large or dense, overcrowding occurs, resulting in increased competition for mates and resources, aggression, parasite transmission, and sometimes cannibalism (6). If the individual is isolated, it poses a challenge to finding mates and resources and induces a chronic stress response (6). Using animal models, the molecular, cellular, and behavioral effects of social isolation can be studied by separating animals of social species from their social group. This has been done using rodents (7,8), bees (9), ducklings (10), zebrafish (11), and fruit flies (12–17). Although *Drosophila* are not eusocial insects, they tend to aggregate and can adapt their behavior and physiology to other flies to function as part of a group (6). In *Drosophila melanogaster*, chronic social isolation has been shown to reduce sleep, increase feeding, and contribute to a transcriptomic profile indicative of a starvation-like brain state (18). However, the cited study only examined the effects of chronic social isolation on activity and feeding in males.

Our goal was to develop a battery of behavioral tests and phenotypes that could be used to assess sex differences in the fly response to social isolation. We identified a set of behaviors that reflected sex-specific changes following seven days of social isolation. The sex differences in these phenotypes prompted us to perform RNA sequencing (RNA-seq) on male and female head samples of socially isolated and group-housed control flies for target identification. Manipulations of one of the emerging target genes, *takeout (to*) suggest mechanistic relationships between isolation, sex, gene expression changes, and sucrose-feeding behavior in the value-based feeding decision (VBFD) test. Overall, this study supports the use of *Drosophila melanogaster* as a valid non-mammalian model for elucidating the neuro-molecular mechanisms of social stress-induced behavioral change.

## Results

### Locomotion increases to a greater extent in males compared to females following social isolation

To characterize social isolation as a valid fly stress model, we performed a series of behavioral tests that could capture changes of potential translational relevance. We recorded fly walking behavior in a 55mm diameter arena and used EthoVision tracking software (Noldus Information Technology Inc., Wageningen, the Netherlands) to quantify distance traveled in a 10-minute period to see if isolation induced hypo- or hyperactivity, and if the behavioral effect differed by sex. We found that social isolation increased distance traveled in both males (Fig 1A, *p* < 0.0001) and females (Fig 1A, *p* < 0.0001), but that the magnitude of the change was significantly greater in males (Fig 1B, *p* < 0.0001).

**Figure 1.**
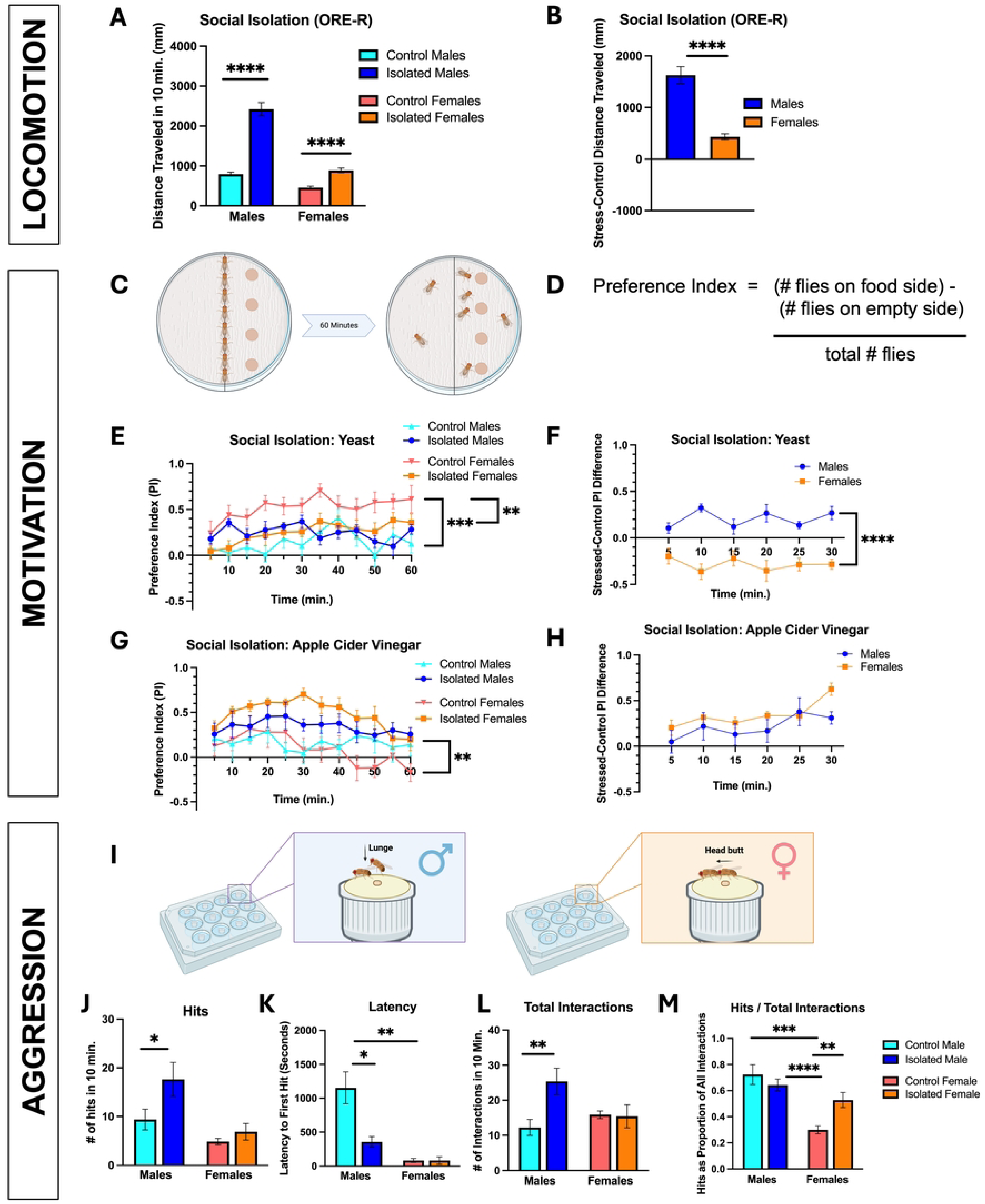
Social isolation differentially alters locomotor, motivated, and aggressive behavior between male and female flies. **A)** Flies that experienced social isolation (n = 99 individuals) traveled greater distances in a 10-minute period than controls (n = 100 individuals) for both males (U = 1729, p < 0.0001) and females (U = 2808, p < 0.0001). **B)** Taking differences in distance traveled between socially isolated flies and the control average for each sex revealed that the increase in locomotion in isolated males relative to control males was significantly greater (n = 99, U = 2850, p < 0.0001) than that observed between isolated females and control females (n = 99). **C)** Set up: flies were briefly anesthetized, positioned along the midline of the arena, and recorded during 60 minutes of exploration time. One half of the arena contained four dots of yeast paste or apple cider vinegar, and the other half had no food. Created with BioRender.com. **D)** A preference index (PI) was calculated at five-minute intervals, where PI = (the number of flies on the food side of the arena) - (the number of flies on the empty side of arena) / the total number of flies. A value of 0 can be interpreted as at chance, due to equal numbers of flies being on each side. **E)** Isolated females (n = 12 groups) are less attracted to yeast paste relative to control females (n = 12 groups, F(1, 19) = 8.65, p = 0.0084). The baseline difference between control males (n=12 groups) and control females (F(1, 18) = 18.43, p = 0.0004) is abrogated upon social isolation. **F)** When comparing the differences in preferences between isolated and control flies, isolated male flies (n = 13 groups) are relatively more attracted to yeast, whereas isolated females are relatively less attracted (n = 11 groups, F(1, 22) = 46.08, p < 0.0001). **G)** Isolated females (n = 12 groups) are more attracted to apple cider vinegar relative to control females (n = 12 groups, F(1, 14) = 14.55, p = 0.0019). **H)** However, trends do not differ significantly between males (n = 8 groups) and females (n = 8 groups) when examining differences in apple cider vinegar preference between isolated and control flies. **I)** Set up: caps filled with standard fly food and topped with yeast paste were placed in a 12-well plate. Same-sex pairs of flies from the same experimental condition were aspirated into each well and recorded for 60 minutes. Male “hits” were defined as lunges and female “hits” were defined as head butts. Created with BioRender.com. **J)** Socially isolated males (n = 16 pairs) hit more compared to group housed controls (n = 16 pairs, U = 76, p < 0.05). There was no difference in the number of hits between isolated (n = 15 pairs) and control females (n = 15 pairs, U = 101, p = 0.87). **K)** The latency between the flies’ first encounter on the food cup and the first hit was lower in socially isolated males relative to controls (U = 71, p < 0.03). Female controls had a significantly shorter latency to first hit compared to male controls (U = 40, p = 0.002) that did not differ from that of isolated females (U = 79, p = 0.57). **L)** Isolated males had a greater number of social interactions, aggressive or non-aggressive, than control males (U = 49.5, p = 0.002) in the 10 minutes following the first hit. **M)** Measuring the proportion of all social interactions that were hits, both control (U = 37, p = 0.0006) and isolated males (U = 14, p < 0.0001) had hits as a greater proportion of all social interactions relative to control females. However, social isolation significantly raised this proportion in females (U = 39.5, p = 0.002) to the levels of males. Error bars are ± SEM. \**P* < 0.05, \*\**p* < 0.01, \*\*\**p* < 0.001, \*\*\*\**p* < 0.0001.

### Social isolation impacts motivation to seek food (yeast) differentially in males versus females

We next decided to assess whether social isolation differentially affected the goal-seeking behavior of males and females, specifically in the context of food motivation. To determine if social isolation alters motivated behavior in flies, we used two different stimuli, yeast paste and apple cider vinegar, to measure flies’ motivation to explore a physical space containing food. Yeast volatiles, either alone or as byproducts of fermenting fruit, attract *Drosophila* (*19–22*). Apple cider vinegar, through its high acetic acid content, mimics the odor of rotting fruit, which attracts flies as a source of food and mates. We developed a novel behavior test, inspired by an existing sucrose preference test (23), to assess whether social isolation alters the motivation of flies to investigate an attractive food or food odorant, and if so, whether sex differences are evident.

In our set up (Fig 1C), briefly anesthetized flies were placed along the midline of a petri dish that was filled with agarose and topped with a piece of parafilm and filter paper, to limit the flies’ space and encourage exploration. Flies were recorded for 60 minutes, and a preference index was calculated (Fig 1D) at 5-minute intervals, to get a sense of the flies’ “preference” over time between a side that had yeast or fermenting fruit odor (apple cider vinegar) and a side that contained nothing.

The preference index (PI) was calculated such that a value of 0 indicated no discrimination between the food side versus non-food side. Using a repeated measures two-way ANOVA, we found that social isolation reduced females’ preference for yeast (Fig 1E, F(1, 19) = 8.65, p = 0.0084) in that it eliminated the difference between males and females observed in the control groups, in which females’ preference had exceeded that of males (Fig 1E, F(1, 18) = 18.43, p = 0.0004). Upon subtracting the control group average from individual PI values of socially isolated males and females at each time point, we found a significant difference between males and females in how each stress group deviated from its control group (Fig 1F, F(1, 22) = 46.08, p < 0.0001). Following social isolation, females demonstrated a reduced preference for the yeast side of the arena relative to control females, while isolated males showed an increased preference for the yeast side compared to control males. In contrast, using apple cider vinegar as an attractive odorant, we observed a significant difference in preference index between control males and control females (Fig 1G, F(1, 14) = 14.55, p = 0.0019) but did not see an effect of social isolation in either sex. For apple cider vinegar, both sexes were more attracted to the odorant side of the arena following social isolation relative to controls, yielding no sex difference (Fig 1H).

### Social isolation increases aggression in males and females across different measures

After testing flies on both hyperactivity and food motivation, we assessed aggression, a behavior that has been previously associated with social isolation in fly models (12,16,24). We used a set up (Fig 1I) similar to one used by other research groups studying *Drosophila* aggression (25): microtube caps were filled with standard fly food, topped with yeast paste, and placed in a 12-well plate, such that each well contained a single cap. This set up created an arena, where flies would encounter each other and begin fighting for territory and food. We defined aggressive “hits” in male flies as the lunge, a behavior in which the aggressor rears up on his hind legs and quickly snaps down on his opponent (25). Aggressive “hits” in females were defined as the head butt, a movement in which the aggressor produces an abdominal thrusting-type motion and strikes the opponent with her head (26). Same-sex pairs of flies were aspirated into a well and recorded for 60 minutes. We noted the time at which the flies first encountered each other on the food cup to measure *latency to first hit* from that initial encounter. A shorter latency to first hit indicates a higher level of aggression (27). Once the first hit occurred, we counted the number of hits and any social interactions that did not result in a hit for the next 10 minutes. We found that social isolation increased the number of hits in males (p < 0.05) but not females (p = 0.87) (Fig 1J). Social isolation also decreased latency to first hit in males (p < 0.03) but not females (0.57), relative to the respective control groups (Fig 1K). Interestingly, females at baseline may have a shorter latency to first hit compared to males. This is reflected in the significant difference in latency to first hit between male controls and female controls (p = 0.002). It is possible that we did not observe a difference between socially isolated females and control females in latency to first hit because the latency in control females is so short to begin with, creating a floor effect. Isolated males also showed a greater number of total social interactions (both aggressive and non-aggressive encounters) compared to controls (p = 0.002) which was not evident in females (Fig 1L). Finally, when examining aggressive hits as a proportion of all social interactions (Fig 1M), there is a difference between isolated females and their controls, such that hits make up a significantly greater proportion of social behaviors in isolated females (p = 0.002). This difference is not seen between isolated males and control males. Both control males (p = 0.0006) and isolated males (p < 0.0001) showed a higher hit proportion compared to control females, but isolated females’ increase in hit proportion rose to the extent that there was no difference between isolated females and isolated males. Thus, the increase in number of hits observed in isolated males may be a product of the fact that isolated males are also interacting more overall; there does not seem to be an increase in the frequency of hitting relative to other social behaviors. Isolated females’ increased proportion of hits out of total social interactions in combination with females’ overall short latency to hit may indicate enhanced female territoriality.

### Social isolation enhances sucrose feeding in females

The value-based feeding decision (VBFD) test was the final test to assess whether social isolation would be reflected in this measure. We followed an existing protocol (28), in which flies were placed in an arena for two hours to feed freely on either a blue solution of sucrose, a more nutritious sugar, or a red solution of arabinose, a less nutritious sugar (Fig 2A). The developers of this behavioral test argue that the results of the VBFD indicate the flies’ cognitive status, as feeding on the more nutritious food source necessitates learning and decision-making, and this discrimination ability declines with age and in models of neurodegenerative disease (28). We decided to interpret the results of this test at face value, as a readout of sugar consumption, to complement the previously described tests of motivated behavior. We used the color (blue, red, purple, or none) of the crop, the fly equivalent of the mammalian stomach, to sort the flies into categories of which sugar they primarily consumed (Fig 2B). We performed additional testing to confirm that the flies did not have a color preference between blue and red.

**Figure 2.**
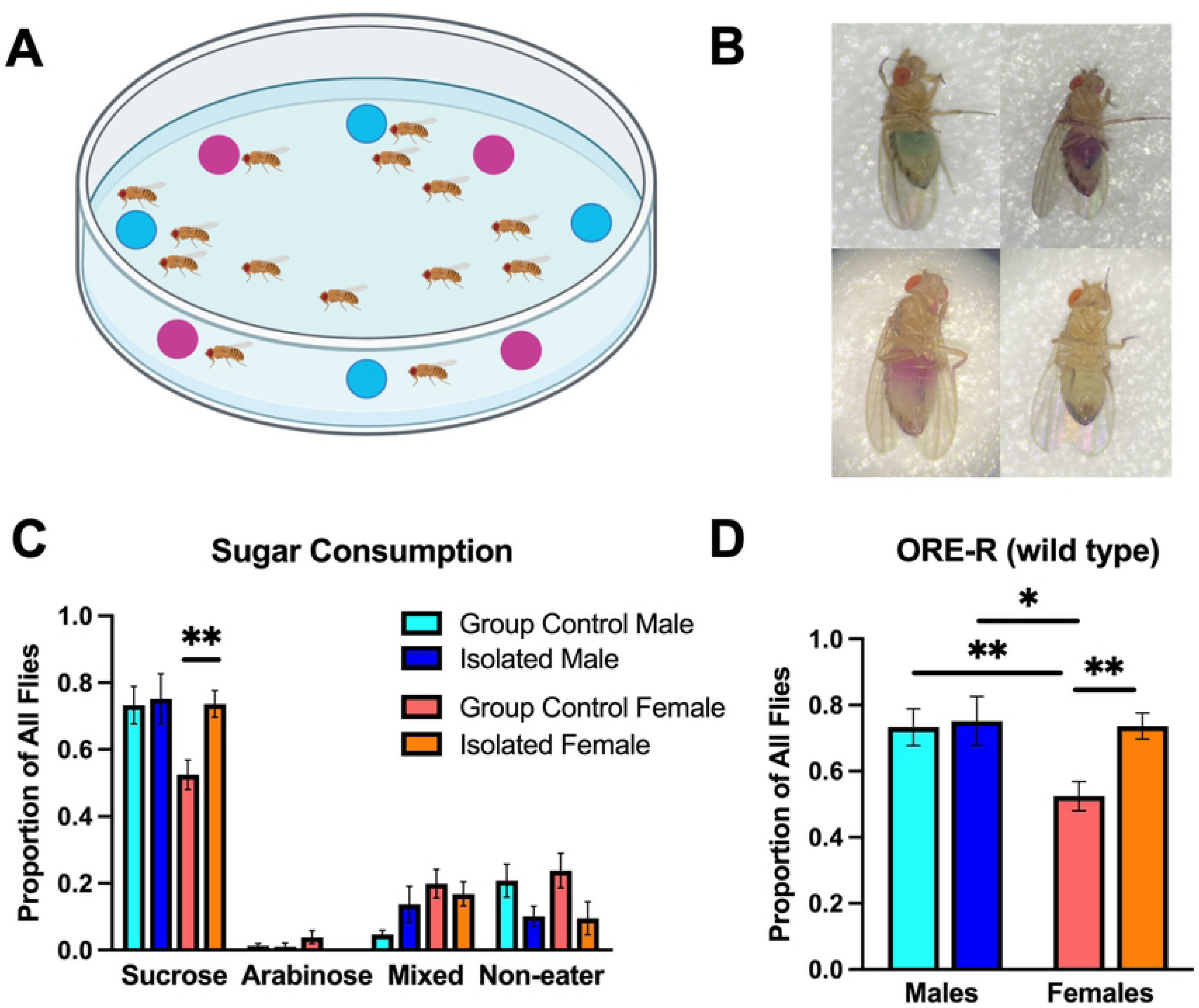
Social isolation increases sucrose consumption in ORE-R females. **A)** Set up: flies were briefly anesthetized and allowed to explore an agarose plate with 2 different sugar solutions for 2 hours. Sucrose (150mM) was dyed blue, and arabinose (150mM) was dyed red. Created with BioRender.com. **B)** Flies were identified (clockwise from top left) as either blue, purple, none, or red and counted at the end of each 2-hour period. All flies shown are female, except for none (bottom right). **C)** Social isolation increased sucrose feeding in females (t(15) = 3.14, p = 0.007). **D)** Control (n = 13 groups, t(22) = 2.85, p = 0.009) and isolated (n = 10 groups, t(19) = 2.67, p = 0.02) males both consume more sucrose than control females (n = 11 groups), but there are no significant differences between these groups and isolated females. Error bars are ± SEM. \**p* < 0.05, \*\**p* < 0.01.

Following social isolation, the proportion of flies consuming sucrose increased significantly in females (Fig 2C, t(15) = 3.14, p = 0.007). No differences were observed in feeding choice between control males and isolated males. Focusing on sucrose, the nutritious sugar used in this test, we observed that both control males (Fig 2D, t(22) = 2.85, p = 0.009) and isolated males (Fig 2D, t(19) = 2.67, p = 0.02) had a greater proportion of flies consuming sucrose relative to control females, but that these differences were erased when comparing males to socially isolated females. Thus, isolation is associated with females increasing their sucrose-feeding behavior to more closely resemble that of males.

### Social isolation uncovers sex differences across behaviors

The outcomes of all behavioral tests are summarized and compared by sex in Table 1. Overall, social isolation produced behavioral alterations across tests of activity, motivation, aggression, and sucrose-feeding. Following isolation, males increased their activity, their motivation to explore a space containing yeast paste or apple cider vinegar odorant, their number of aggressive hits (lunges), and overall social interactions in the aggression assay compared to male controls. Males’ latency to strike an opponent decreased, indicating heightened aggression. Females also increased their activity, motivation to explore the apple cider vinegar odorant, percentage of overall social interactions that were aggressive hits (head butts), and sucrose consumption relative to arabinose and non-feeding. Isolated females decreased their exploration of the yeast paste side of the arena relative to control females.

**Table 1.**
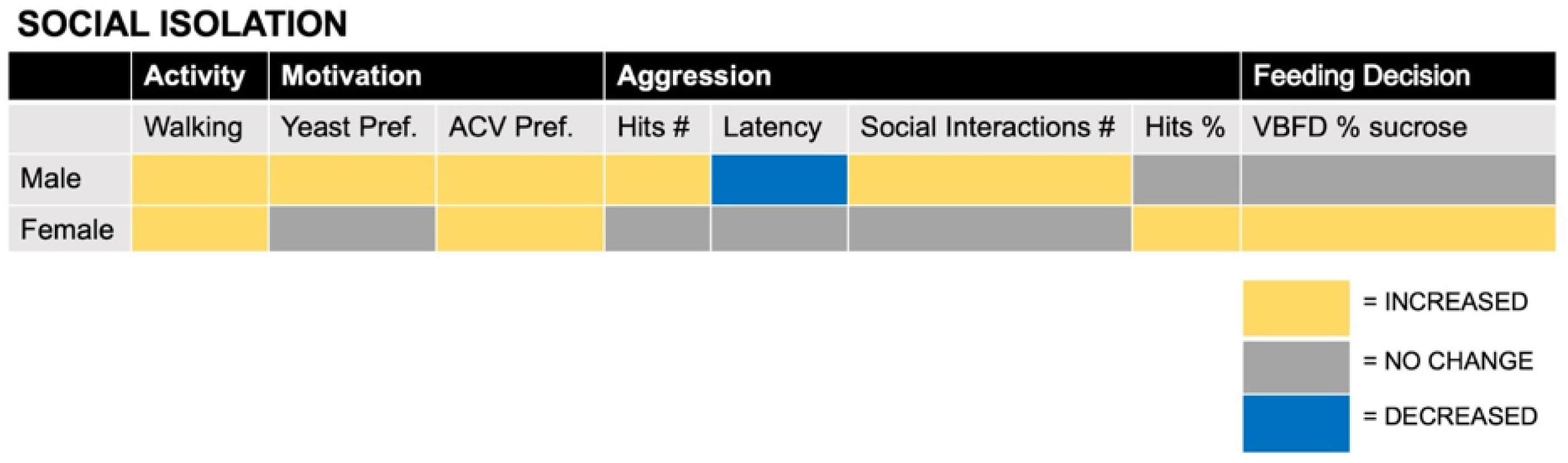
Summary of behavioral changes and sex differences following social isolation.

### Social isolation does not impact lifespan

To further explore the stress effects of social isolation, we examined whether isolation affected longevity. Newly-eclosed flies were set up in groups of 50. The “stressed” group experienced social isolation for seven days and then returned to group-housed standard conditions for the remainder of the experiment. Flies were checked every 24 hours for new deaths until no flies remained alive. We found no differences in survival between socially isolated and control flies for males (S1 Fig, χ² = 2.71, df = 1, p = 0.10) or females (S1 Fig, χ² = 0.25, df = 1, p = 0.62). Additional statistical tests, beyond the log-rank test, are detailed in Table 2.

**Table 2.** Statistical analysis of survival for social isolation.

| MALES | Chi square | df | P value | FEMALES | Chi square | df | P value |
| --- | --- | --- | --- | --- | --- | --- | --- |
| Log-rank test | 2.711 | 1 | 0.0997 | Log-rank test | 0.2481 | 1 | 0.6184 |
| Gehan-Breslow-Wilcoxon test | 2.448 | 1 | 0.1177 | Gehan-Breslow-Wilcoxon test | 0.006334 | 1 | 0.9366 |
| Hazard ratio (Mantel-Haenszel): 0.6770 (0.4255, 1.077) |  |  |  | Hazard ratio (Mantel-Haenszel): 1.114 (0.7276, 1.707) |  |  |  |
| Hazard ratio (logrank): 0.7560 (0.5089, 1.123) |  |  |  | Hazard ratio (logrank): 1.102 (0.7355, 1.651) |  |  |  |

### Differential expression analysis indicates female-specific overexpression of fat body-synthesized proteins following social isolation

RNA samples were extracted in triplicate from whole fly heads for each experimental group: social isolation (SI) male, control male, SI female, and control female. The samples were then submitted to the Princeton Genomics Core Facility for cDNA library preparation and sequencing (see Methods > Transcriptome analysis by RNA Sequencing (RNA-Seq) > Sequencing). Upon receiving the FASTQ files, we processed these raw files through a bioinformatics pipeline to identify differentially expressed genes (see Methods > Transcriptome analysis by RNA Sequencing (RNA-Seq) > Analysis pipeline).

We began by generating a Multidimensional Scaling (MDS) Plot as a dimensionality reduction approach to discern the overall similarity and dissimilarity across our samples (S2 Fig). Of our triplicates, we removed one sample from each of the female groups that failed this quality test for more accurate target discovery. The resulting MDS plot shows a separation between the samples, along the top two dimensions that account for the greatest percentage of variance in log fold change, by sex and condition.

Next, we used *voom* to prepare the data for linear modeling with *limma*. Our design matrix modeled the relationship between gene expression and condition (SI female, control female, SI male, and control male) adjusted for batch. Specifically, we focused on identifying differences in gene expression between the sexes under social isolation, relative to their respective controls. Differentially expressed genes were filtered based on an FDR-adjusted p- value of p_adj_ < 0.05 to determine significant genes with positive or negative changes when comparing socially isolated males to control males, socially isolated females to control females, and the interaction between social isolation and sex. The heatmap in Fig 3A depicts the genes that met these criteria when comparing the interaction between social isolation and sex. We observed clusters of genes that were upregulated in isolated females compared to control females and males (Fig 3A). Upon running these significant genes through Metascape (29), a gene annotation and pathway analysis resource, we identified changes in groups of genes related to cell division (e.g. mitotic cell cycle process, asymmetric cell division) and metabolism (e.g. cellular catabolic process, fatty acid metabolic process) (Fig 3B).

**Figure 3.**
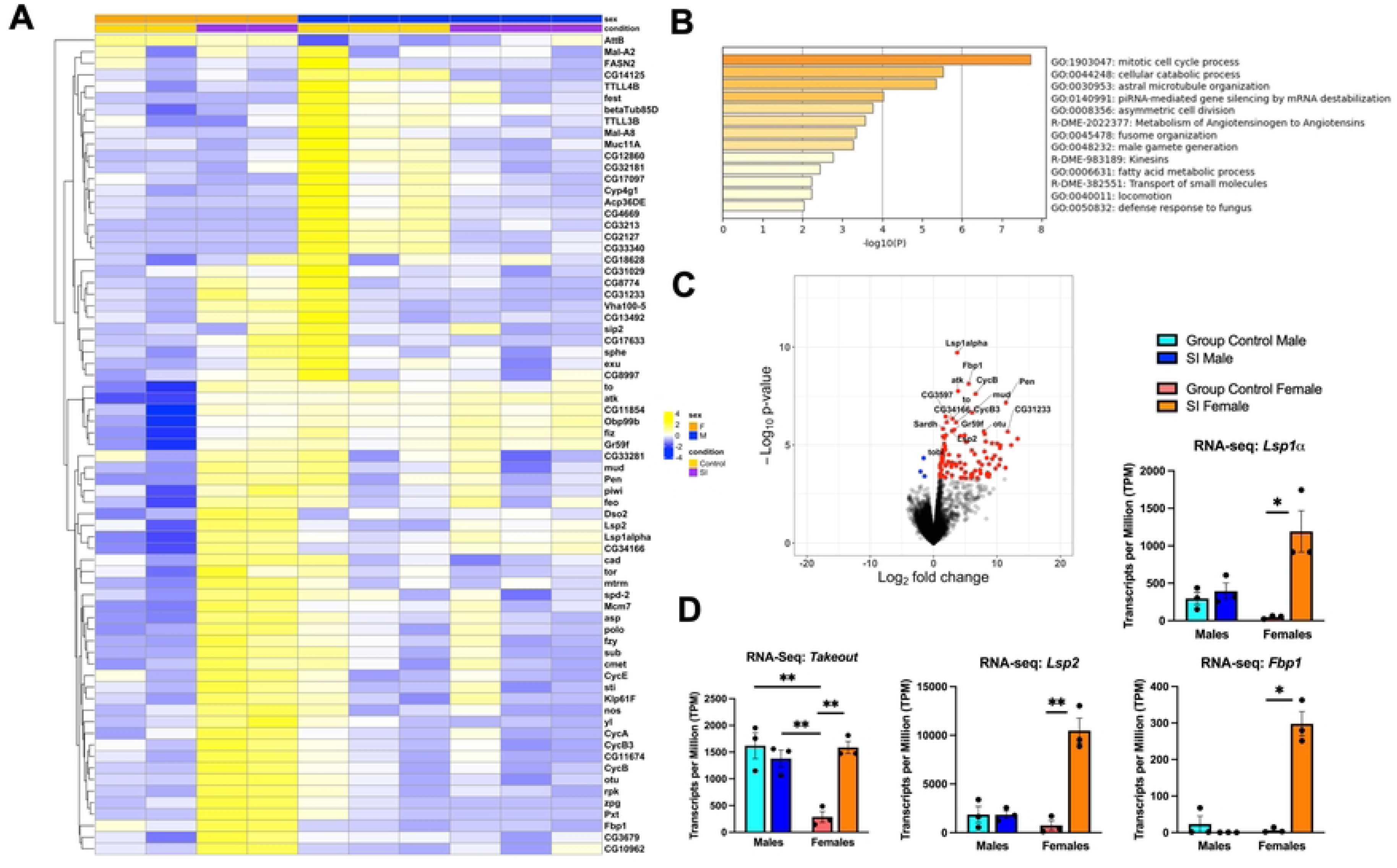
Socially isolated females demonstrate upregulation of multiple genes related to metabolic processes relative to control females and males. **A)** Heat map depicting the scaled (z-score transformed) log2 counts per million of genes differentially expressed (absolute value of log2-fold change > 2 and adjusted p-value < 0.05) when comparing the difference between SI female and control female with the difference between SI males and control males. **B)** Gene ontology (Metascape) of the significantly differentially expressed genes captured in the heat map reflects changes in broad categories of cell division and metabolism. **C)** The top differentially expressed genes (absolute value of log2-fold change > 1 and adjusted p-value < 0.05), when comparing the interaction between social isolation and sex, reflect upregulation in isolated females. The red points represent the genes upregulated in socially isolated females (relative to female controls) compared with socially isolated males (relative to male controls). In contrast, blue points represent the genes (*CG11211*, *VhaM9.7-a*, and *AttB*) upregulated in socially isolated males compared to the socially isolated females, relative to their respective controls. **D)** Gene expression, quantified in transcripts per million (TPM) from RNA-seq, indicates elevation in *takeout, Lsp2*, *Fbp1,* and *Lsp1α* in isolated females relative to control females that is not seen in males. At baseline, control females had lower TPMs compared to control males (t(4) = 5.07, p = 0.007) or isolated males (t(4) = 5.79, p = 0.004). However, socially isolated females had significantly higher TPMs compared to control females (t(4) = 8.56, p = 0.001) which are comparable to those of males. TPMs were also elevated following social isolation in females for *Lsp2* (t(4) = 7.06, p = 0.002), *Fbp1* (t(2.06) = 8.64, p = 0.01 with Welch’s correction), and *Lsp1α* (t(4) = 4.14, p = 0.01). Closed circles represent values derived from individual RNA samples, which were extracted from 200 fly whole heads. Error bars are ± SEM. \**p* < 0.05, \*\**p* < 0.01.

Additionally, entering the significantly differentially expressed genes (comparing the interaction of social isolation and sex) into the STRING database (30) returned a network map of potential functional protein-protein interactions among them (S3 Fig). Applying the Markov Cluster (MCL) Algorithm (31) to our data then parsed our genes into 21 potential functional clusters, many of which coincided with the Metascape gene ontology results. Clusters with the strongest functional enrichment included genes related to mitotic cell cycle (S3 Fig), female reproduction (S3 Fig), and hemocyanins/hexamerins and secreted storage proteins (S3 Fig). STRING calculates measures of enrichment strength, signal, and statistical significance. *Strength* captures the size of the enrichment effect by calculating log_10_(observed/expected), where the (observed/expected) ratio represents the ratio between the number of genes in our network annotated with a term and the number that we would expect to see annotated with this term in a random network of the same size. *Signal* measures a weighted harmonic mean between the (observed/expected) ratio and -log(false discovery rate). *False discovery rate* (FDR) describes the significance of the enrichment, with p-values corrected for multiple testing with the Benjamini-Hochberg procedure (32).

We also used STRING to visualize our top functional enrichments, based on gene count, signal, and FDR, categorized by biological process, local network clustering, and *Drosophila* phenotype (S4 Fig). Reiterating the findings from our clustering, the top biological process terms related to mitotic cell cycle processes and cell division (S4 Fig). Local network clustering showed enrichment in egg activation, piRNA metabolic process, and hemocyanins/hexamerins (S4 Fig). Finally, our significant genes were implicated in *Drosophila* phenotypes of abnormal meiosis and female sterility (S4 Fig).

Finally, we used our significance cut-off of p_adj_ < 0.05 to visualize our differentially expressed genes in a volcano plot (Fig 3C). Only three significant genes were downregulated when comparing the interaction of social isolation and sex, while 105 were upregulated. The most significantly differentially expressed genes, by adjusted p-value, are listed in Table 3, along with their log fold change values. These genes were all upregulated in isolated females compared to isolated males, when taking into account the differences between the isolated groups and their respective controls.

**Table 3.**
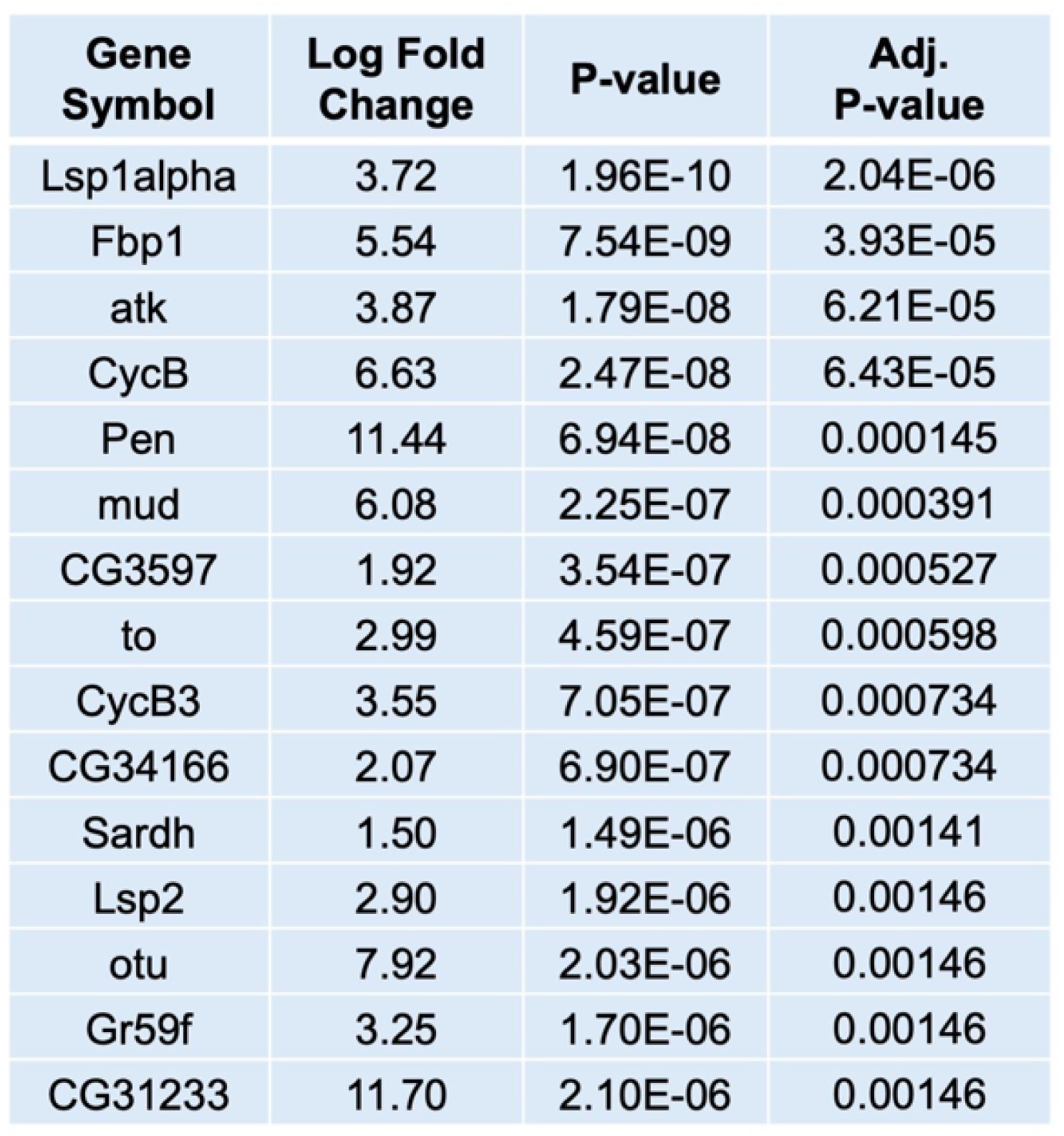
Top differentially expressed genes (interaction of social isolation and sex).

| Gene Symbol | Log Fold Change | P-value | Adj. P-value |
| --- | --- | --- | --- |
| Lsp1alpha | 3.72 | 1.96E-10 | 2.04E-06 |
| Fbp1 | 5.54 | 7.54E-09 | 3.93E-05 |
| atk | 3.87 | 1.79E-08 | 6.21E-05 |
| CycB | 6.63 | 2.47E-08 | 6.43E-05 |
| Pen | 11.44 | 6.94E-08 | 0.000145 |
| mud | 6.08 | 2.25E-07 | 0.000391 |
| CG3597 | 1.92 | 3.54E-07 | 0.000527 |
| to | 2.99 | 4.59E-07 | 0.000598 |
| CycB3 | 3.55 | 7.05E-07 | 0.000734 |
| CG34166 | 2.07 | 6.90E-07 | 0.000734 |
| Sardh | 1.50 | 1.49E-06 | 0.00141 |
| Lsp2 | 2.90 | 1.92E-06 | 0.00146 |
| otu | 7.92 | 2.03E-06 | 0.00146 |
| Gr59f | 3.25 | 1.70E-06 | 0.00146 |
| CG31233 | 11.70 | 2.10E-06 | 0.00146 |

An analysis of expression quantity data (transcripts per million, TPM), of some of the top genes identified by our model, confirmed isolated female specific overexpression of RNA transcripts for *larval serum protein-2* (*Lsp-2*), *larval serum protein-1ɑ* (*Lsp-1ɑ*), *fat body protein 1* (*Fbp1*), and *takeout* (*to*). *Lsp-2, Lsp-1ɑ, Fbp1*, and *to* are all synthesized in fat body cells (33–36), with the first three genes functioning as amino acid storage proteins and *to* being implicated in feeding behavior. *To* expression is higher in control (Fig 3D, t(4) = 5.07, p = 0.007) and isolated males (Fig 3D, t(4) = 5.79, p = 0.004) relative to control females, but this male-female difference is abolished when females have undergone social isolation. *To* expression is significantly higher in isolated females compared to control females (Fig 3D, t(4) = 8.56, p = 0.001). Similarly, social isolation increases *Lsp-2* (Fig 3D, t(4) = 7.06, p = 0.002)*, Lsp-1ɑ* (Fig 3D, t(4) = 4.14, p = 0.01), and *Fbp1* (Fig 3D, t(2.06) = 8.64, p = 0.01) expression relative to control levels in females but not in males.

Insect larval serum proteins, or hexamerins, are generated in the pericerebral fat body, the functional analog of both the liver and fat tissue in vertebrates (37–39). The fat body plays a role in storing excess energy in the form of triglycerides and glycogen (40). It also generates hexamerin storage proteins that are accumulated during the insect’s larval stage (41,42) and secreted from the fat body into the hemolymph. Hexamerins are then circulated as reservoirs of amino acids and metal ions that can be accessed during periods of high metabolic need, such as metamorphosis and reproduction (42). This is beneficial for insects such as flies, which undergo a period in their life cycle when they cannot access food (in the pupal stage). Amino acid storage consequently fluctuates over developmental time periods to ensure proper development and adult fitness (43). The fat body also plays a crucial role in nutrient sensing by communicating the nutritional status of the organism to the brain via multiple molecular signaling pathways. These signals activate insulin-producing cells (IPCs), which form and release *Drosophila* insulin-like peptides (DILPs) to regulate levels of glucose circulating in hemolymph (44).

*Lsp-2* encodes a major hexamerin that serves as a nutrient store in the late larval and pupal stages but also continues accumulation in adulthood (36). Recently, Fbp1 has also been characterized to play a role in the reuptake of LSPs and FBPs from the hemolymph back into fat body cells (43). Based on the results from our STRING functional enrichment, social isolation may disrupt pathways associated with reproduction. If amino acid stores are accessed during reproduction, in response to high metabolic demand, it follows that the stores will accumulate during periods when reproduction is not occurring.

*To* was originally characterized as a gene encoding a secretory protein that is mainly expressed in the fly head (45) and mediates circadian feeding behavior (46). It is also expressed in organs related to feeding and olfaction and is upregulated in these structures in response to starvation, with *to* mutants dying shortly following the onset of starvation (45). *To* expression may also be regulated by sex-specific factors. In the pericerebral fat body surrounding the brain, the male-specific forms of Doublesex (Dsx^M^) and Fruitless (Fru^M^) proteins activate *to* expression, but the female-specific form of Doublesex (Dsx^F^) protein suppresses it (35). This is consistent with our *to* TPM data from the male and female controls (Fig 3D). Moreover, *to* has been associated with male courtship behavior, which significantly decreases upon feminization of fat body cells via expression of the female Transformer protein (Tra^F^) in males (35,47). To the best of our knowledge, the effect of social isolation on *to* expression has not been studied.

To confirm the RNA-seq results, we used qPCR as a secondary method to validate the expression of *to* in male and female fly heads. Using remaining RNA from the same samples that were submitted for sequencing, we analyzed expression of *to* in SI males and SI females relative to their respective controls. The qPCR data largely corroborated the RNA-seq findings. Overexpression of *to* following social isolation was specific to females and differed significantly from the gene expression change between isolated and control males (S5 Fig, t(4) = 6.14, p = 0.004).

### qPCR of ORE-R brains suggests that social isolation increases expression of *takeout* in females but not males

Following RNA-seq profiling and validation, we identified *to* as a gene demonstrating female-specific overexpression in the head following social isolation. However, this finding raised the question of whether *to* expression was changing in the brain or elsewhere in the head. For instance, the sex difference in isolation-induced *to* expression could stem from transcriptional changes in the pericerebral fat body, where *to* has been established to be transcriptionally regulated in a sexually dimorphic manner, or in the antennae (35) or foregut (48). To gain a better understanding of whether *to* expression is changing within or outside of the fly brain, the brains of additional male and female flies that had experienced either social isolation or group housing (control conditions) were isolated, and RNA was extracted. The subsequent qPCR experiment again reflected female-specific overexpression of *to* following isolation (S6 Fig, t(10) = 2.28, p < 0.05).

### Adult *Drosophila* express *to* primarily in olfactory receptor neurons, epithelial cells of the gastrointestinal tract, and antennal nerve glia

In addition to validating our bulk RNA-seq results with qPCR, we also compared our findings with existing data from *Drosophila melanogaster* RNA-seq atlases. RNA-seq data from the modENCODE project (49) indicate that adult male *Drosophila* have higher whole organism-wide expression of *to* compared to adult females, which is consistent with our data comparing males to control female heads and brains. Based on data from the Fly Cell Atlas project (50), *to* was most highly expressed in (sensory) neurons, sensory organ cells, glial cells, and epithelial cells out of all adult cell types in the *Drosophila* body (S7 Fig). The prominence of *to* across many olfactory receptor neuron (ORN) subtypes (S7 Fig) suggests a potential function for *to* in regulating feeding through sensitivity to food odor. ORNs express chemoreceptors that bind odor molecules, and their axons synapse with projection neuron dendrites in antennal lobe glomeruli. Olfactory information is then transmitted through the circuit to higher order brain regions, the lateral horn and mushroom bodies (51).

The RNA-seq atlas data also showed *to* to be expressed in multiple epithelial cell types throughout the digestive system (S7 Fig), including the cells of the crop, midgut, esophagus, hindgut, and enteroendocrine cells, which secrete hormones to regulate digestion, metabolic homeostasis, and feeding behavior (52). Finally, in glial cells, *to* showed the most widespread expression in adult antennal nerve glia (S7 Fig). Antennal glial cells ensheath ORNs and may modulate the neuronal response to olfactory stimuli (53). It is plausible that sex differences in *to* expression in any of these cell types could contribute to the differential response in sucrose consumption and food motivation between males and females following isolation stress. However, the replication of our whole head RNA-seq finding in isolated brains narrowed our focus to neurons and glia. Because glial cells only constitute approximately 10% of cells in the *Drosophila* nervous system (54), we decided to manipulate *to* expression using GAL4 driver lines specific to *to*-expressing cells and postmitotic neurons.

### Knocking down *to* in *to*-expressing cells decreases sucrose-feeding choice in socially isolated females and increases it in socially isolated males

A complete summary of our manipulations and behavioral findings is provided in Table 4. In order to manipulate the expression of *to* in *to*-expressing cells, we mated virgin female flies with GAL4 expressed under the control of a *to*-specific promoter (*to*-Gal4) to males with a UAS enhancer 1) inducing the generation of RNA interference (RNAi) specifically targeted to *to* mRNA (UAS-*to*^RNAi^), or 2) driving expression of *to* (UAS-*to*). We performed qPCR tests to validate that *to* expression was downregulated in *to*-Gal4xUAS-*to*^RNAi^ flies relative to ORE-Rs and upregulated in *to*-Gal4xUAS-*to* flies relative to ORE-Rs across experimental groups (S13 Fig). The *to*-Gal4xUAS-*to*^RNAi^ offspring were tested in the VBFD test to assess the effect of *to* knockdown in *to*-expressing cells on the proportion of flies that chose to feed on sucrose, a more nutritious sugar, over arabinose, a less nutritious sugar. The “proportion of all flies” dependent measure represents the proportion of all flies in a test group that fed exclusively or almost exclusively on sucrose, compared to all other combinations of feeding choice (a sucrose-arabinose mixture, arabinose only, or neither). Of these, sucrose is the optimal choice for nutritional value.

**Table 4.** Summary of behavioral results in VBFD test across genetic manipulations and experimental groups.

| Effect of manipulation on sucrose feeding (yellow = up, grey = no change, blue = down) |  |  |  |  |
| --- | --- | --- | --- | --- |
|  | Control Males | Social Isolation Males | Control Females | Social Isolation Females |
| *to*<sup>RNAi</sup> in *to*-expressing cells (to-Gal4 x UAS-*to*<sup>RNAi</sup>) | grey | yellow | grey | blue |
| *to* overexpression in *to*-expressing cells (to-Gal4 x UAS-*to*) | grey | grey | grey | blue |
| *to*<sup>RNAi</sup> in neurons (elav-Gal4 x UAS-*to*<sup>RNAi</sup>) | grey | grey | grey | grey |
| *to* overexpression in neurons (elav-Gal4 x UAS-*to*) | yellow | yellow | grey | grey |
| *tra*<sup>RNAi</sup> in neurons (elav-Gal4 x UAS-*tra*<sup>RNAi</sup>) | grey | grey | grey | grey |

Control males, isolated males, control females, and isolated females of *to*-Gal4xUAS- *to*^RNAi^ and each parent genotype were tested. We observed an increase in sucrose feeding in isolated males compared to control males (S8 Fig, t(14) = 2.31, p = 0.04), but a decrease in sucrose feeding in isolated females relative to female controls (S8 Fig, t(15) = 5.35, p < 0.0001). This male isolation-induced increase in sucrose choice is specific to the knockdown manipulation and is not evident across either of the parent genotypes or the wild type. Moreover, knocking down *to* (*to*-Gal4 x UAS-*to*^RNAi^) reverses the trend observed in isolated females relative to control females that is also observed in the UAS-*to*^RNAi^ and ORE-R flies.

Upon examining the effect of genotype on sucrose consumption, within a specific experimental group, we found that *to*-Gal4 x UAS-*to*^RNAi^ isolated females had a significantly *lower* proportion of sucrose-feeding flies compared to all other genotypes: ORE-R (Fig 4B, t(12) = 5.24, p = 0.0002), *to*-Gal4 (Fig 4B, t(13) = 5.53, p < 0.0001), UAS-*to*^RNAi^ (Fig 4B, t(14) = 3.15, p = 0.007), and *to*-Gal4 x UAS-mCherry^RNAi^ (Fig 4B, t(12) = 3.84, p = 0.002). In isolated males, the *to*-gal4 x UAS-*to*^RNAi^ flies increased their proportion of sucrose feeding relative to both parent genotypes (*to*-Gal4: Fig 4D, t(15) = 2.39, p = 0.03; UAS-*to*^RNAi^: Fig 4D, t(14) = 3.84, p = 0.002) and the ORE-Rs (Fig 4D, t(16) = 2.14, p < 0.05).

**Figure 4.**
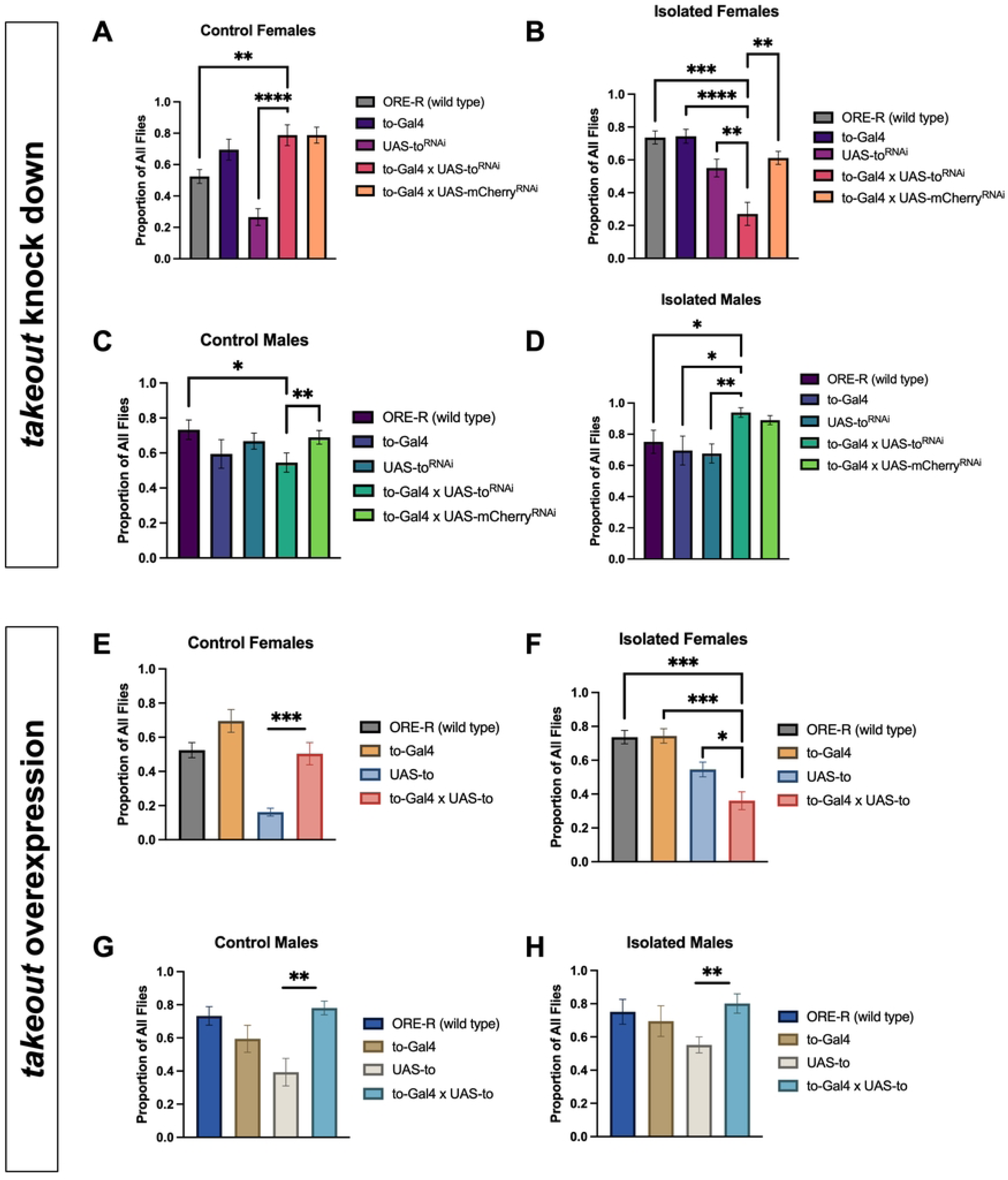
The proportion of all flies that primarily consumed sucrose changes in opposite directions, relative to other genotypes, between isolated females and isolated males upon knock down of *to* in *to*-expressing cells. The proportion of all flies that primarily consumed sucrose decreases relative to other genotypes in isolated females upon overexpression of *to* in *to*-expressing cells. **A)** Sucrose feeding is elevated in control females of the to-gal4 x UAS-to^RNAi^ cross relative to ORE-R (t(18) = 3.41, p = 0.003) and UAS-to^RNAi^ (t(22) = 6.07, p < 0.0001) control females but does not differ from to-Gal4 or to-Gal4 x UAS-mCherry^RNAi^ control females. **B)** Sucrose feeding is reduced in isolated females of the to-gal4 x UAS-to^RNAi^ cross relative to isolated females of all other genotypes: ORE-R (t(12) = 5.24, p = 0.0002), to-Gal4 (t(13) = 5.53, p < 0.0001), UAS-to^RNAi^ (t(14) = 3.15, p = 0.007), and to-Gal4 x UAS-mCherry^RNAi^ (t(12) = 3.84, p = 0.002). **C)** Sucrose feeding is reduced in control males of the to-gal4 x UAS-to^RNAi^ cross relative to ORE-R control males (t(20) = 2.31, p = 0.03) and to-Gal4 x UAS-mCherry^RNAi^ control males (t(14) = 3.89, p = 0.002) but does not differ from parent genotype control males. **D)** Sucrose feeding increases in isolated males of the to-gal4 x UAS-to^RNAi^ cross relative to wild type and parent genotype isolated males: ORE-R (t(16) = 2.14, p < 0.05), to-Gal4 (t(15) = 2.39, p = 0.03), and UAS-to^RNAi^ (t(14) = 3.84, p = 0.002). **E)** Sucrose feeding is elevated in control females of the to-gal4 x UAS-to cross relative to UAS-to control females (t(16) = 4.50, p = 0.0004) but does not differ from ORE-R or to-Gal4 control females. **F)** Sucrose feeding is reduced in isolated females of the to-gal4 x UAS-to cross relative to isolated females of the wild type and parent genotypes: ORE-R (t(11) = 5.54, p = 0.0002), to-Gal4 (t(12) = 5.65, p = 0.0001), and UAS-to (t(13) = 2.74, p = 0.02). **G)** Sucrose feeding is enhanced in control males of the to-gal4 x UAS-to cross relative to UAS-to control males (t(18) = 3.91, p = 0.001) but does not differ from wild type or to-Gal4 control males. **H)** Sucrose feeding is elevated in isolated males of the to-gal4 x UAS-to cross relative to UAS-to isolated males (t(14) = 3.31, p = 0.005) but does not differ from wild type or to-Gal4 control males. \**p* < 0.05, \*\**p* < 0.01, \*\*\**p* < 0.001, \*\*\*\**p* < 0.0001.

### Overexpressing *to* in *to*-expressing cells decreases sucrose feeding in isolated females, but not males, relative to the ORE-R and parent genotypes

Upon testing the *to-*Gal4 x UAS-*to* flies on the VBFD test, we found that isolation (relative to group housing) did not seem to affect sucrose choice in either sex (males: S9 Fig, t(14) = 0.30, p = 0.77; females: S9 Fig, t(15) = 1.59, p = 0.13). However, in isolated females, sucrose feeding choice was depressed in *to*-Gal4 x UAS-*to* flies compared to ORE-Rs (Fig 4F, t(11) = 5.54, p = 0.0002) and both parent genotypes (*to*-Gal4: Fig 4F, t(12) = 5.65, p = 0.0001; UAS-*to*: Fig 4F, t(13) = 2.74, p = 0.02). In both control (Fig 4G, t(18) = 3.91, p = 0.001) and isolated males (Fig 4H, t(14) = 3.31, p = 0.005), sucrose feeding choice was increased in *to*-Gal4 x UAS-*to* flies compared to UAS-*to* males but did not differ from *to-*Gal4 or ORE-R flies of the same experimental condition.

### Driving *to* expression in neurons increases sucrose-feeding in males

After observing sex differences in sucrose feeding upon *to* manipulation in all *to*- expressing cells (using the *to*-Gal4 driver), we also wanted to down- and upregulate expression of *to* in neurons alone. Given that the qPCR results from extracted brains corroborated female-specific upregulation of *to* following chronic social isolation, and the single cell RNA-seq atlas data indicated *to* present across several olfactory receptor neurons, we selected neurons as a logical target to probe whether changes in *to* expression in the brain itself are responsible for sex differences in feeding behavior.

We began by mating *elav*-Gal4 virgin female flies with UAS-*to*^RNAi^ males to generate flies expressing RNAi targeted to *to* in postmitotic neurons. The elav-Gal4 line is commonly used in *Drosophila* research to drive the expression of GAL4 under control of the *elav* gene, which is active in postmitotic neurons throughout development and is expressed throughout most cells of the fly nervous system (55,56). Via qPCR, we validated that *to* expression was downregulated in *elav*-Gal4 x UAS-*to*^RNAi^ flies relative to ORE-Rs across experimental groups (S13 Fig).

We did not observe any differences in sucrose-feeding following social isolation in male or female offspring of the elav-Gal4 x UAS-*to*^RNAi^ cross (S10 Fig). An examination of within-experimental group effects of genotype on sucrose feeding choice also failed to show an effect of *to* knock down in neurons on behavior. Although elav-Gal4 x UAS-*to*^RNAi^ control females increased sucrose feeding compared to ORE-R (Fig 5A, t(16) = 2.25, p = 0.04) and UAS-*to*^RNAi^ (Fig 5A, t(20) = 4.72, p = 0.0001) control females, there was no difference relative to the other parent genotype, *elav*-Gal4. Across isolated females (Fig 5B), control males (Fig 5C), and isolated males (Fig 5D), the elav-Gal4 x UAS-*to*^RNAi^ flies did not differ in their feeding behavior from any other experimental control group (the wild type and parent genotypes) in a statistically significant manner.

**Figure 5.**
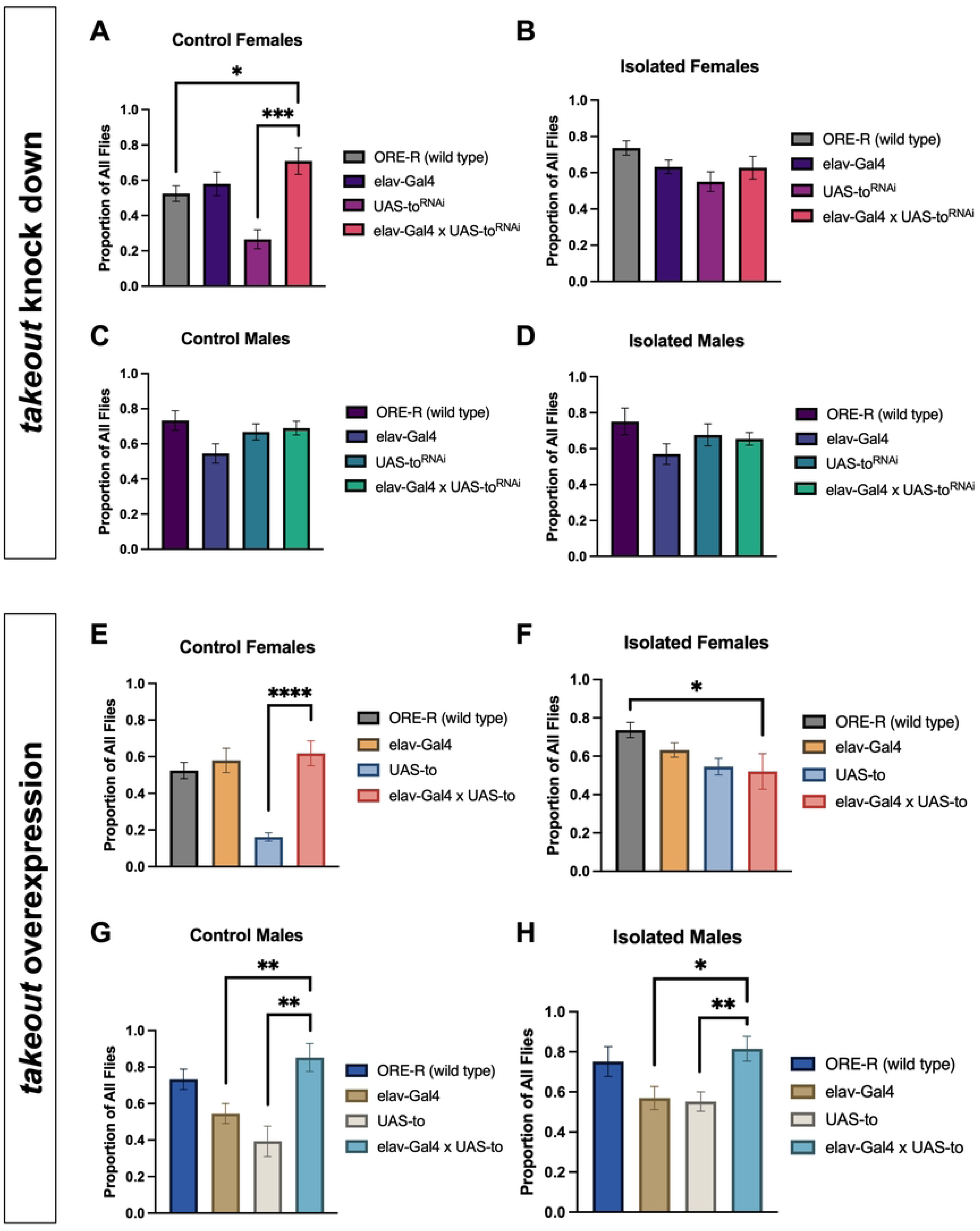
The proportion of all flies that primarily consumed sucrose does not change in any experimental groups upon knock down of *to* in neurons. The proportion of all flies that primarily consumed sucrose increases relative to parent genotypes in control males and isolated males upon overexpression of *to* in neurons. **A)** Sucrose feeding is elevated in control females of the elav-Gal4 x UAS-to^RNAi^ cross relative to ORE-R (t(16) = 2.25, p = 0.04) and UAS-to^RNAi^ (t(20) = 4.72, p = 0.0001) control females but does not differ from elav-Gal4 control females. **B)** Sucrose feeding does not differ between elav-Gal4 x UAS-to^RNAi^, wild type, and parent genotype isolated females. **C)** Sucrose feeding does not differ between elav-Gal4 x UAS- to^RNAi^, wild type, and parent genotype control males. **D)** Sucrose feeding does not differ between elav-Gal4 x UAS-to^RNAi^, wild type, and parent genotype isolated males. **E)** Sucrose feeding is elevated in control females of the elav-gal4 x UAS-to cross relative to UAS-to control females (t(12) = 7.15, p < 0.0001). **F)** Sucrose feeding is reduced in isolated females of the elav-gal4 x UAS-to cross relative to ORE-R isolated females (t(9) = 2.30, p < 0.05). **G)** Sucrose feeding is enhanced in control males of the elav-gal4 x UAS-to cross relative to UAS-to control males (t(15) = 3.63, p = 0.003) and elav-Gal4 control males (t(13) = 3.33, p = 0.005). **H)** Sucrose feeding is also elevated in isolated males of the elav-gal4 x UAS-to cross relative to UAS-to isolated males (t(12) = 3.30, p = 0.006) and elav-Gal4 isolated males (t(12) = 2.71, p = 0.02). \**p* < 0.05, \*\**p* < 0.01, \*\*\**p* < 0.001, \*\*\*\**p* < 0.0001.

Next, we set up elav-Gal4 x UAS-*to* crosses to assess the behavioral effect of *to* overexpression in neurons.When we tested the elav-Gal4 x UAS-*to* flies, there was not an effect of isolation on sucrose choice in females (S11 Fig, t(9) = 0.87, p = 0.41). For within-experimental group effects of genotype on sucrose feeding choice, we noted that in males, both control flies (Fig 5G) and socially isolated flies (Fig 5H) increased sucrose-feeding relative to both parent genotypes. Thus, overexpressing *to* in postmitotic neurons seems to increase sucrose-feeding in males, but not females, irrespective of social housing or isolation.

### Masculinizing female neurons via RNAi targeted to *transformer (tra)* does not alter sucrose feeding choice

Finally, we wanted to test whether altering the sexual identity of neurons to make female neurons more male-like affected sucrose feeding choice. To do this, we crossed elav-Gal4 virgin females with UAS-*transformer* (*tra*)^RNAi^ males. This genetic manipulation has been used previously as a tool to masculinize neurons in female flies (57). The UAS-*tra*^RNAi^ line uses double-stranded RNA to knock down *tra*, a gene in the *Drosophila* sex determination pathway that is crucial for female somatic sexual differentiation (58). In female flies only, the gene *Sex-lethal* (*Sxl*) produces a functional protein, which then splices *tra* as a downstream target. This yields a functional tra protein in females but not males, which then performs sex-specific alternative mRNA splicing on *doublesex* (*dsx*) and *fruitless* (*fru*) (59–63). Sex-specific forms of *dsx* and male *fru* then act as transcription factors to influence sex-specific development of somatic and neural tissues and behavior (64–66). Hence, we were interested in knocking down *tra* in neurons, to assess whether this manipulation made sucrose feeding choice more “male-like” in females. Using qPCR, we validated that *tra* expression was downregulated in *elav*-Gal4 x UAS-*tra*^RNAi^ flies relative to ORE-Rs across experimental groups (S13 Fig).

Overall, we did not observe changes in sucrose-feeding in isolated males and isolated females relative to their respective controls in the elav-Gal4 x UAS-tra^RNAi^ offspring (S12 Fig, males: t(17) = 1.42, p = 0.17; females: t(17) = 0.91, p = 0.38). Within-condition comparisons across genotypes also did not reflect any clear differences in sucrose feeding choice upon *tra* knock down in neurons. We did not see any differences across all genotypes in control females (Fig 6A) or isolated males (Fig 6D). Interestingly, qPCR validation revealed that knockdown of *tra* resulted in decreased expression of *to* across all experimental groups relative to ORE-R levels (S13 Fig). Since knocking down *to* expression in neurons through the elav-Gal4 x UAS-to^RNAi^ manipulation did not yield any changes in sucrose-feeding, it is consistent that decreasing *to* expression through neuron masculinization similarly did not produce clear changes in this behavior.

**Figure 6.**
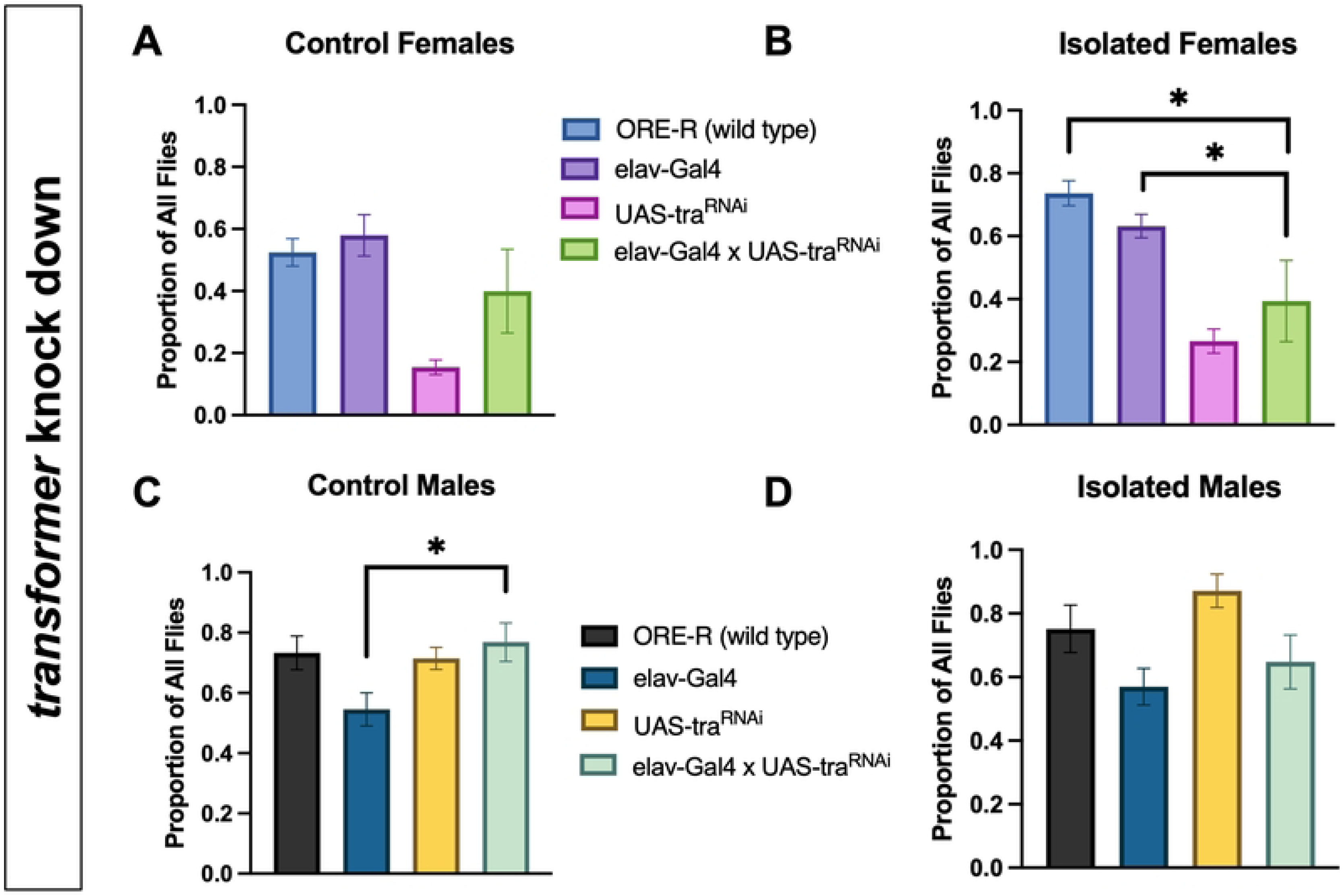
The proportion of all flies that primarily consumed sucrose does not differ between elav-Gal4 x UAS-tra^RNAi^ flies and other genotypes, across experimental groups. **A)** No significant differences were detected in sucrose feeding between elav-gal4 x UAS-tra^RNAi^ control females and control females of the wild type and parent genotypes. **B)** Sucrose feeding is reduced in isolated females of the elav-gal4 x UAS-tra^RNAi^ cross relative to ORE-R (t(10) = 2.54, p = 0.03) and elav-Gal4 isolated females (t(14) = 2.19, p < 0.05). **C)** Sucrose feeding is elevated in elav-gal4 x UAS-tra^RNAi^ control males relative to elav-Gal4 control males (t(13) = 2.61, p = 0.02). **D)** No significant differences were detected in sucrose feeding between elav-gal4 x UAS-tra^RNAi^ isolated males and isolated males of the wild type and parent genotypes. \**p* < 0.05.

Hence, through these genetic manipulations, we found that either up- or downregulating *to* in *to-*expressing cells decreases sucrose-feeding in socially isolated females. However, these same manipulations produce either increases or no changes in sucrose-feeding in socially isolated males. Altering gene expression of *to* in neurons does not seem to generate changes in sucrose-feeding in females, but *to* overexpression in neurons may increase sucrose-feeding in males, regardless of social condition.

## Discussion

### Summary

We first aimed to establish social isolation as a valid stressor in flies, using behavioral and survival data to capture stress-induced phenotypic changes. Overall, we found changes in males and females across behavioral tests of locomotion, food motivation, and aggression following seven days of social isolation. There were no significant effects of social isolation on lifespan in either males or females.

In the behavioral data, we also observed a potential female shift in macronutrient preference from protein to sugar following social isolation, implicated by isolated females’ decreased exploration of the “yeast side” of the arena, increased exploration when yeast was replaced by apple cider vinegar, and increased preference for sucrose. Isolated males were less discerning, showing increased exploration for both yeast and apple cider vinegar and no change in sucrose consumption compared to controls. It could be evolutionarily advantageous for females to prioritize sugar over protein consumption while under the environmental stress of isolation, to favor survival over reproduction. Although yeast is composed of sugar, protein, and salt, flies rely on yeast as their primary protein source, and both males and females increase their consumption of yeast prior to mating to prepare for reproduction (67). Mated females also increase their yeast consumption for the benefit of their offspring to the detriment of their own stress resistance and lifespan, suggesting a fitness-fecundity trade off (67). While social isolation may increase general feeding in males, based on the literature, the male macronutrient preference may be less clear, given that males do not have the reproductive demand of egg-laying. Additionally, the different manifestations of isolation-induced aggression between males and females may reflect different ways in which males versus females display territoriality over a food source.

We did not observe any differences in lifespan between socially isolated flies and group-housed controls of either sex. Clearly, social isolation was sufficiently stressful to alter behavior but insufficiently stressful to decrease lifespan. We isolated the flies for seven days and then returned them to group housing for the remainder of their lives. It is possible that the isolation experience fostered resilience in the flies once they were returned to standard conditions, and that we may have observed declines in lifespan had we employed lifelong social isolation. Another explanation could be that all experimental groups were same-sex. Long term same-sex pairing has been found to decrease lifespan in males and females (68), so it is possible that our group-housed controls were not completely unstressed controls, making it more difficult to detect stressed versus control differences. It would be worthwhile in the future to compare the survival of isolated males and females to that of a mixed-sex group-housed control group, to rule out a confounding stress effect of same-sex group living.

We next focused on the sex-specific effects of social isolation on gene expression, and the potential stress-signaling pathways in which these genes participate. To understand how social isolation impacts gene expression, and subsequently behavior, differentially in male versus female flies, we used RNA-seq as an unbiased approach for target identification. Using STRING, we identified functional enrichments suggesting that gene expression may be altered in socially isolated females compared to isolated males, relative to their respective controls, in a manner that disrupts reproduction. Hemocyanin and hexamerin proteins were also upregulated in isolated females. Hemocyanins largely play a role in oxygen transport but have also been associated with innate immunity, wound healing, hormone transport, and ecdysis (moulting) (69). Hexamerins act as storage proteins in the hemolymph of insects that provide the organism with amino acids and energy during non-feeding periods. However, hexamerins may also play roles in cuticle formation, hormone transport, and humoral immunity (41). Finally, our differential expression analysis revealed that the gene *to* is upregulated in females, but not males, following isolation. We validated this RNA-seq result with qPCR of both the original samples and new samples derived from extracted brains.

Upon manipulating *to* expression with various GAL4-UAS combinations, we found that knocking down *to* in *to*-expressing cells decreases sucrose-feeding in isolated females relative to control females of the same genotype but increases sucrose-feeding in isolated males relative to same-genotype control males. Within-condition comparisons of behavior by genotype supported these findings. For isolated females, *to*-Gal4 x UAS-*to*^RNAi^ flies showed decreases in sucrose consumption compared to ORE-R (wild type) flies, flies of each parent genotype, and *to*-Gal4 x UAS-*mCherry*^RNAi^ flies (an inert RNAi control). For isolated males, *to*-Gal4 x UAS-*to*^RNAi^ flies showed increases in sucrose consumption compared to ORE-Rs and flies of each parent genotype.

In the model that aimed to overexpress *to* in *to*-expressing cells, we did not see an effect of condition (social isolation) on sucrose choice in either sex for the *to*-Gal4 x UAS-*to* flies. However, when examined by experimental condition and sex, there was an effect of genotype in isolated females such that *to*-Gal4 x UAS-*to* isolated females consumed less sucrose relative to ORE-R and parent genotype isolated females. Taken together, these results suggest that *takeout* may function in isolated females like a neuromodulator; when either downregulated or upregulated in *to*-expressing cells, in the context of social isolation, sucrose feeding choice is suppressed. The opposite behavioral response was observed in males. Isolated males increased sucrose consumption upon *to* knock down and maintained their level of consumption when *to* was overexpressed in *to*-expressing cells.

When *to* expression was manipulated in neurons, we observed no effects of *to* knock-down. However, *to* overexpression increased sucrose-feeding exclusively in males regardless of isolation or control status. It should be noted that qPCR validation of the overexpression model was inconclusive (S13 Fig). Masculinizing neurons in females via *tra* knock down did not produce differences in feeding behavior in either control or isolated flies relative to other genotypes.

Based on these results, the overexpression of *to* driving sucrose-feeding in isolated females may be located in non-neuronal cells or tissues within the fly head. It is possible that our qPCR of extracted brains detected *to* upregulation in non-neuronal cell types, or that cells from the pericerebral fat body remained on the brains when they were extracted. *to* was originally characterized as a pericerebral fat body gene that is upregulated in response to starvation (45). This would be consistent with social isolation inducing a starvation-like brain state (18), thus upregulating *to* expression. The single-cell RNA-seq atlas data indicate that *to* is also widely expressed in ORNs, epithelial cells, and glia. ORNs play an important role in feeding behavior by enabling odor-detection during foraging. Although it does not seem that *to* expression in neurons is the source or key driver of female isolation-induced sucrose preference, *to* could be expressed in glia that support the function of these neurons.

In the future, it may be worthwhile to try masculinizing the fat body instead of neurons, to see if masculinizing this tissue in control females increases sucrose-feeding in the VBFD test to male-like levels. One study has suggested that feminizing the fat body in adult male *Drosophila* decreases both courtship behavior and the levels of takeout protein in the head (47). As many as 46 sex-biased genes have been identified in the fly head, with some being localized to the fat cells (70). These sex-specific transcripts in the head fat body may encode protein products, which are secreted and act on the brain to generate sex differences in behavior. Moreover, manipulations of *tra* and its cofactor *tra-2* in the context of starvation or oxidative stress have been shown to reverse the expected male versus female stress response, measured by heart rate and locomotor patterns (71). These results suggest that effectors produced in the fat body may influence sexually dimorphic actions in the brain outside of its own transcriptome.

Alternatively, *to* could influence behavior through the binding of juvenile hormone, a fly hormone that canonically regulates developmental transitions. The protein encoded by *to* is highly structurally similar to juvenile hormone binding protein (JHBP), and it contains a putative binding domain for juvenile hormone (45,46). The secretion of takeout into the hemolymph from the pericerebral fat body and its binding of juvenile hormone could either hinder juvenile hormone signaling, by reducing levels of the hormone actively circulating, or facilitate juvenile hormone acting on its targets by functioning as a carrier protein.

The question remains that if social isolation induces a starvation-like brain state (18), why are changes in *to* expression and sucrose feeding female-specific? One explanation could be a behavioral ceiling effect in males, given that wild type control and socially isolated males both consume more sucrose than control females at baseline. However, the sex differences in *to* expression and feeding behavior could also be connected to changes in sexual behavior and reproduction in response to chronic stress. We found evidence of a sex difference in motivated behavior in the yeast preference test in addition to the VBFD test. Following social isolation, fewer females yet more males were observed on the yeast side of the arena relative to their respective controls. Isolated females’ decreased exploratory behavior around yeast and increased consumption of a nutritious sugar (sucrose) could be interpreted as a shift in macronutrient preference to prioritize survival over reproduction.

In the present study, it could be that in a stressful environment, in which the fly is perceiving social isolation similarly to hunger, there could be a macronutrient shift from protein to sugar in females, as the females attempt to bolster their own stress resistance and lifespan. Females may have a lower stress threshold than males to start changing feeding and mating behavior, as males do not have to support and lay eggs. Food-deprived male flies do tend to prioritize feeding over courtship, and this preference can be reversed on the time scale of minutes by consuming protein-rich food (72). This male behavioral switch from feeding to courtship occurs when ingested amino acids trigger the release of Diuretic hormone 31 (Dh31), a neuropeptide hormone, from the enteroendocrine cells in the gut into the hemolymph. Dh31 then binds to its receptors in the brain in two distinct neuronal populations: one that inhibits feeding, and one that stimulates courtship (72). It would be interesting to add courtship and sexual receptivity to our battery of behavioral tests to test the hypothesis further that social isolation-induced *to* upregulation in females functions to divert nutritional resources away from reproduction and towards survival. It might follow that females may be less sexually receptive following social isolation, if a stressful environment is not conducive to reproduction.

There is some initial evidence that takeout may play a role in the tradeoff-switch between fertility and fitness, and that it may exert its effects by binding juvenile hormone, thus restricting juvenile hormone’s signaling activity. Overexpression of *to*, specifically in the fat body, has been found to increase fly longevity and decrease male courtship behavior. These effects of *to* overexpression are nullified when flies are treated with methoprene, a juvenile hormone analog (73). If *to* overexpression improves lifespan, this could also explain the lack of a survival difference between isolated females and group-housed control females in the present work. Female isolation-induced increases in *to* expression could have improved lifespan, counteracting any negative stress effects of social isolation that shorten lifespan. Finally, in addition to overexpression, downregulation of *to* via a loss-of-function mutation also decreases male courtship behavior (35). These researchers hypothesized that takeout may play a redundant role with other Fru^M^-regulated factors that influence mating in males. Since there is evidence of both upregulation and downregulation of *to* reducing male courtship, this strengthens the argument that takeout may act similarly to other neuromodulators.

### Limitations and future directions

The present study was limited with respect to certain aspects of our behavioral testing and RNA sequencing. In the VBFD test, we quantified sucrose feeding choice by categorizing the flies by crop color as primarily consuming sucrose, arabinose, both sugars (in roughly equal quantity), or nothing, and then calculating a proportion of sucrose-consuming flies out of all animals. We did not quantify the exact amount of food consumed, nor more closely examine how flies were making their choice. For example, for flies that consumed both sugars, we did not distinguish between flies that began eating arabinose and then switched to sucrose midway through the feeding period, or flies that were indiscriminately feeding between sucrose and arabinose during the entirety of the experiment. We could use an approach such as the CApillary FEeder (CAFE) assay (74), which directly measures food intake in *Drosophila* via capillary tube liquid feeding, to better quantify actual food consumption in the future.

Our RNA-seq analyses were based on a low number of replicates, which necessitated validation using qPCR tests. We also used bulk RNA-seq, whereas single-cell RNA sequencing (scRNA-seq) would have been more informative in determining if female-specific *to* overexpression is driven by a specific cell type.

A final limitation in this work was that our manipulations of *to* expression only focused on spatial control of gene expression. In our *to*–Gal4 x UAS-*to*^RNAi^ flies, we knocked down *to* in all *to*-expressing cells of the body, not just the ones in the head that were detected in our RNA profiling. For future experiments, it would be helpful to perform immunohistochemistry or fluorescence in situ hybridization to determine where *to* is being overexpressed in the isolated female head or brain and then target manipulations to a more specific cell or tissue type. We could also expand our genetic tools to allow for temporal control to see if activation or inactivation of to+ cells during the behavior experiment impacts feeding choice. This would only be relevant to sex differences in *to* expression in a specific population of neurons, but optogenetic manipulations could extend our studies to elucidate the role of *to*-expressing neurons and their potential targets in the neural circuitry underlying sex differences in food choice within a chronic stress context.

## Conclusion

Overall, chronic social isolation alters internal states and behaviors across animal species, and it may do so in a manner that differs between males and females. We have found that social isolation in *Drosophila* impacts macronutrient preference and sucrose consumption differentially on the basis of sex, and that these behavioral changes may depend on sex differences in the expression of the gene *to* in the fly’s head. By changing the expression of storage proteins and proteins such as takeout in response to stress and hunger, flies can adjust their behavior to achieve energetic balance. Chylomicrons, low-density lipoproteins, and amino acid binding proteins in humans bear some functional similarity to these molecules, in that they are involved in the response to feeding and nutritional depletion. Collectively, our findings support the use of *Drosophila melanogaster* as a model system for exploring female-specific molecular mechanisms related to stress-induced changes in feeding, metabolism, and possibly fertility.

## Materials and methods

### Fly stocks and rearing

All fly stocks were raised on standard medium (cornmeal/yeast/molasses/agar) at 25℃ and 40% relative humidity on a fixed 12-h light/dark cycle. Male and female *Oregon-R* flies (wild type) and lines from the Bloomington *Drosophila* Stock Center (BDSC) were used for the majority of these experiments. Behavioral experiments were conducted within 8 hours after lights-on. The following fly lines were obtained from the BDSC: *to-Gal4* (#80938), *elav-Gal4* (#8765), *UAS-to* (#81000), and *UAS-to*^RNAi^ (#55982). *Oregon-R* and *UAS-mCherry*^RNAi^ were gifts from Dr. Jodi Schottenfeld-Roames, Princeton University. The *UAS-tra*^RNAi^ (Vienna *Drosophila* Resource Center #2560, also known as traIR) stock was kindly provided by Dr.

Rachel Monyak, Stonehill College.

### Experimental setups and design

#### Social isolation

For the social isolation stress model, we used an experimental procedure for chronic social isolation already characterized in the literature (18). Newly-eclosed male and female flies were housed together for three to five days to acquire social experience. Flies were then housed either singly or in same-sex groups of 25 for seven days. Within 24 hours of the final day of isolation or group housing, flies were either frozen on dry ice for RNA extraction, tested for behavior, or moved to/kept in same-sex group housing for survival analysis.

#### Survival

To measure survival, flies (n = 50 per experimental group) underwent social isolation or group-housed control conditions for seven days, following the previously described protocol. At the end of the stress period, all flies were moved into group housing in standard food vials. The flies were counted once per day, and any deaths were recorded, along with the date of death, until all flies were deceased.

### Behavioral tests

#### Locomotion assay

Flies were transferred to the testing chamber via aspiration or brief anesthesia with CO_2_. Upon analysis, we found that the use of CO_2_ did not alter activity when compared to aspiration. Each chamber consisted of a 55mm diameter petri dish filled with 1% agarose gel, to provide the flies with enough room to walk but not fly. The flies were put into same-sex groups of 10, recorded for 10 minutes, and frozen on dry ice. Videos were analyzed using EthoVision XT video tracking software (Noldus Information Technology, VA).

#### Motivated behavior tests

Flies were tested in the arena previously described in “locomotion assay” to evaluate anhedonia-like behavior or discrimination of a food source. Flies were briefly anesthetized using CO_2_ and lined up along the midline of a filter paper evenly divided into two halves: one with yeast paste and one with nothing. Behavior was recorded for 60 minutes, and a preference index (PI) was calculated at five minute intervals as PI = [(# of flies on the yeast side) – (# of flies on the empty side)] / (total). This test was repeated using apple cider vinegar in place of yeast paste, as an additional appealing food stimulus.

#### Aggression assay

To test aggression, we modified an experimental protocol previously described by the Kravitz lab (25). Testing occurred within 24 hours following the end of social isolation or crowding stress. Twelve-well plates were used as behavior chambers for testing same-sex, same-experimental condition pairs of flies. For group housed controls, flies in a fighting pair were taken from different vials to account for social novelty. To create an arena/territory, white plastic microtube caps were filled with standard fly food, topped with a dot of yeast paste, and placed in the wells. A transparent plastic covering with drilled holes was placed on top of the well plate, so that flies could be gently aspirated into the wells at the same time. The holes were then plugged with short screws. Fights were recorded for one hour using a Sony Digital Handycam camcorder (Sony Group Corporation, Tokyo, Japan).

Only behavior that took place in the arena was scored. Flies that did not encounter each other on the food cup within 45 minutes of recording were excluded from analysis. A goal sample size of n = 15 fighting pairs was decided prior to the start of experiments. For the fighting pairs, the times of the first encounter on the food cup and the first hit were recorded to calculate latency to initiate fighting. Fights were then scored for 10 minutes following the first hit (lunge for males or head butt for females). The male lunge was defined as a fly rearing up on its hind legs and snapping downwards onto its opponent. The female head butt was defined as a fly thrusting forward horizontally and then recoiling from the opponent. Number of hits, latency, number of overall encounters, and hits as a percentage of all encounters were used as measures to quantify and interpret aggressive behavior.

#### Value-based feeding decision (VBFD) test

The value-based feeding decision (VBFD) test (28) was used as a two-choice assay to examine feeding decision making. Flies were briefly anesthetized with CO_2_ and placed in the center of the arena used for the locomotion and motivated behavior tests, a 55 mm diameter petri dish filled with 1% agarose gel and topped with a layer of parafilm. Sugars were dissolved in plain water to make two solutions: a 150 mM sucrose solution and a 150 mM arabinose solution. Each solution was labeled with a blue dye (0.01% erioglaucine disodium salt, Acros Organics, Cat# 229730250) or red dye (0.1% Food Red No. 106, TCI, Cat# F0143). Alternating droplets (20 μL volume) of each liquid food were placed evenly around the perimeter of the dish, such that there were eight droplets total, four of each solution. Flies were allowed to explore the arena and feed in the dark for two hours. At the end of this period, flies were collected and counted under a microscope. They were scored by color: blue (ingested primarily blue food), red (ingested primarily red food), purple (ingested both foods), or non-eater (no color accumulation in the abdomen).

### RNA extraction

For each sample, flies were decapitated on dry ice. RNA was extracted and purified from homogenized whole heads using a phenol chloroform organic extraction. Samples were also treated using either the RQ1 RNase-Free DNase kit (Promega Corporation) or the TURBO DNA-*free*™ Kit (Invitrogen) to address genomic DNA contamination. Total RNA samples were sequenced by the Princeton Genomics Core or used for downstream applications. The samples submitted for RNA-seq were extracted from 200 whole heads. To obtain brains, 20-40 *Drosophila* heads were dissected one at a time in cold Schneider’s medium. Brains were transferred to a microtube on ice containing 200 μL of Schneider’s medium, briefly spun down on a benchtop centrifuge, and then frozen at −80°C. Prior to homogenization in TRIzol (Invitrogen), samples were spun down, and the Schneider’s medium covering the brains was removed.

### Quantitative Polymerase Chain Reaction (qPCR)

RNA was reverse transcribed to complementary DNA (cDNA) using the SuperScript™ III First-Strand Synthesis System (Invitrogen). Random primers and a 200 ng total RNA template were used in a 20 μL reaction, following the instructions.

For real-time qPCRs, samples used SYBR Green PCR Master Mix (Qiagen or Applied Biosystems, Thermo Fisher Scientific). The PCR was performed in a 10 μL reaction including 3 μL of cDNA and 7 μL of master mix. The master mix for each gene consisted of 0.5 μL of each primer (10 μM), 5 μL SYBR Green, and 1 μL RNase-free water. We used the following PCR conditions: initial incubation at 95°C for 15 min, followed by 40 cycles of 95°C for 15 s and 60°C for 1 min. Relative quantification was performed using the 2^−△△CT^ method (75).

### Transcriptome analysis by RNA Sequencing (RNA-Seq)

#### Sequencing

RNA-seq was performed by the Princeton University Genomics Core Facility. The integrity of total RNA samples was assessed on the 2100 Bioanalyzer system using the RNA 6000 Pico Chip (Agilent Technologies, CA). The poly-A containing RNA transcripts were enriched from one microgram of total RNA for each sample using oligo-dT beads, further fragmented, and converted to cDNA and Illumina sequencing libraries using the PrepX RNA-seq library preparation protocol on the Apollo 324^TM^ NGS Library Prep System (Takara Bio, CA). Different DNA barcodes were attached to each library. The RNA-seq libraries were examined on Agilent Bioanalyzer DNA High Sensitivity chips for size distribution, quantified by Qubit fluorometer (Invitrogen, CA), and pooled at equal molar amounts. Each library pool was denatured and sequenced on one Illumina NovaSeq 6000 S Prime lane using the 100 cycle v1.5 kit. Raw sequencing reads were filtered by Illumina NovaSeq Control software, and only the Pass-Filter (PF) reads were used for further analysis.

#### Analysis pipeline

Sequences were aligned to the *Drosophila melanogaster* genome (BDGP.46) (76,77) using STAR (v. 2.7.10b) (78). Duplicates were removed using the Picard Toolkit (v. 2.27.5) (79) MarkDuplicates function. The number of reads mapping to each gene was determined using htseq-count (HTSeq v. 2.0.5) (80). Differential gene expression analysis was performed in R (v. 4.4.0). All read counts from rRNA and pseudo-rRNA genes were removed. Normalization was performed using edgeR (v. 4.4.1) (81). Multidimensional scaling (MDS) plots were used to identify outlier samples, which were then removed from the analysis. Data was prepared for linear modeling with limma (v. 3.60.6) (82) using voom (83). The design matrix was constructed to model the relationship between gene expression and condition (Female-SI, Female-Control, Male-SI, Male-Control) adjusted for batch. Contrasts were used to compare gene expression differences between the following groups: 1. SI and control females, 2. SI and control males, 3. SI vs controls ((SI Female + SI Male)/(Control Female + Control Male). A fourth contrast was included to compare the difference between SI Female and Control Female with the difference between SI Male and Control Male. Differentially expressed genes were filtered based on an FDR-adjusted p-value of < 0.05 to determine significant genes with positive or negative values of log2FoldChange when comparing socially isolated males to control males, socially isolated females to control females, and the interaction between social isolation and sex. Heatmaps and additional plots were created using the R packages ComplexHeatmap (v. 2.21.2) (84) and ggplot2 (v. 3.5.1) (85).

### Statistical analysis

Statistical analyses were performed using GraphPad Prism 10.4.0 software (GraphPad Software, Inc., La Jolla, CA). The Shapiro-Wilk test of normality (passed at *p* > 0.05) was used to determine whether data followed a parametric distribution. Unpaired t-tests were used for parametric data assuming equal variances, and Welch’s t-tests were used if the samples had unequal variances based on the result of the F-test. If the data did not pass the test for normality distribution, nonparametric testing was used. All nonparametric data were analyzed with the Mann–Whitney *U* test for between-group comparisons. Effects were deemed statistically significant at *p* < 0.05. The Grubbs’ test was applied to identify and exclude outliers in the data. Log-rank tests were used to determine differences in survival curves, with *p* < 0.05 deemed statistically significant. A repeated measures 2-way ANOVA was used to compare differences between stressed and control flies’ preferences for yeast or apple cider vinegar between males and females over time. All statistics for two-way (S1 Table) and one-way (S2 Table) ANOVAs are provided in Supporting Information.

## Acknowledgments

We would like to acknowledge Gillian Hilscher, Sarah Brown, and Ella DePaolo for their contributions to the development and execution of the behavioral tests. We thank Dr. Caroline Palavicino-Maggio (McLean Hospital/Harvard Medical School) for guidance regarding the aggression testing.

## Supporting information

**S1 Fig. Social isolation (SI) does not affect lifespan in male or female flies.** Log-rank tests showed no significant differences in survival between isolated males (n = 50) and group-housed control males (n = 50, χ² = 2.71, df = 1, p = 0.10) or isolated females (n = 50) and group-housed control females (n = 50, χ² = 0.25, df = 1, p = 0.62).

**S2 Fig. Multidimensional Scaling reflects similarity and dissimilarity within and between experimental groups.** One sample (that was most dissimilar from the other two) was removed from each of the female groups to improve target discovery. Key: open = control, solid = social isolation; blue = male, orange = female; square = batch 1, circle = batch 2.

**S3 Fig. Significant differentially expressed genes, when comparing the interaction between social isolation and sex, are connected in broader networks of protein interactions. A)** All connected interactions are depicted, with non-connected genes removed. We specified a setting of “medium interaction strength” in STRING. The thickness of the connecting line between genes roughly corresponds to the strength of data support for gene interaction. **B)** One of our clusters that showed the strongest functional enrichment included genes implicated in the mitotic cell cycle process (strength = 0.82, signal = 0.78, FDR: p = 0.0003). **C)** Another top cluster consisted of genes connected to piwi, with functional enrichments in cellular process involved in reproduction in multicellular organism (strength = 0.49, signal = 0.45, FDR: p = 0.006), piRNA metabolic process (strength = 1.31, signal = 1.72, FDR: p = 6.97e-08), and egg activation and regulation of oskar mRNA translation (strength = 1.60, signal = 1.62, FDR: p = 9.30e-07). **D)** The third cluster of interest contains three genes encoding secreted (strength = 0.76, signal = 0.83, FDR: p = 8.01e-05) storage proteins (strength = 1.89, signal = 0.70, FDR: p = 0.004): Lsp2, Lsp1alpha, and Obp99b. All four genes are categorized as hemocyanin/hexamerin (strength = 1.72, signal = 0.92, FDR: p = 0.0006).

**S4 Fig. Significant differentially expressed genes, when comparing the interaction between social isolation and sex, are enriched across biological process, local network clustering, and *Drosophila phenotype* terms relating to cell division, reproduction, and hemocyanin/hexamerin proteins. A)** Top biological process terms include mitotic cell cycle process and nuclear cell division. **B)** Top local network cluster terms include egg activation, piRNA metabolic process, and regulation of mitotic cell cycle phase transition. **C)** Top *Drosophila* phenotypes include abnormal meiotic cell cycle and female sterility.

**S5 Fig. Validation of the RNA-seq-derived target genes with qPCR indicates enhanced expression of *takeout* in socially isolated females relative to isolated males. A)** A comparison of relative expression fold change values reveals that *takeout* is overexpressed in females but not males following social isolation. The difference in expression between isolated and control flies is significantly greater in females versus males (t(4) = 6.14, p = 0.004). **B)** Social isolation enhances *Lsp-2* expression in males and females (t(4) = 2.56, p = 0.06). **C)** Social isolation enhances *Lsp-1α* expression in males and females (t(4) = 2.21, p = 0.09). **D)** We did not find a statistically significant male-female difference in expression of *Fbp1* following social isolation (t(4) = 1.71, p = 0.16). Data points represent individual cDNA samples for a particular condition relative to a control average. Error bars are ± SEM. \*\**p* < 0.01.

**S6 Fig. *Takeout* expression is elevated in the brains of females but not males following social isolation.** Using relative expression fold change as a measure of the magnitude of change in *takeout* (*to*) expression between the social isolation and control groups, we observed that the values for males are randomly distributed around no change (1), while the female values are largely positive and significantly higher compared to males (t(10) = 2.28, p < 0.05). Data points represent individual cDNA samples for a particular condition relative to a control average. Error bars are ± SEM. \**p* < 0.05.

**S7 Fig. Single cell RNA-seq data from the Fly Cell Atlas identify *takeout* (*to*) expression in neuronal, glial, and epithelial cell types. A)** Colors in the ribbon indicate that *to* expression was identified in that cell type through single nucleus RNA sequencing (snRNA-seq) performed by the Fly Cell Atlas project^36^. A redder color on the blue-red gradient indicates that a greater proportion of cells of that particular cell type express *to*. White means that there is no available snRNA-seq data available for that cell type. **B)** For neurons, *to* was expressed primarily in adult olfactory receptor neurons. **C)** For epithelial cells, *to* was expressed across numerous cell types of the gastrointestinal tract (crop, midgut, esophagus, and hindgut). **D)** For glial cells, *to* expression was most widespread within the glia of the adult antennal nerve.

**S8 Fig. Knock down of *to* via RNA interference (RNAi) in *to*-expressing cells significantly decreases sucrose feeding choice in isolated females and increases sucrose feeding in isolated males relative to their respective controls. A)** In ORE-R flies, our preliminary results indicated a female-specific increase in sucrose feeding following social isolation (t(15) = 3.14, p = 0.007). **B)** Flies expressing GAL4 under the control of a *to*-specific promoter do not show differences in sucrose feeding following social isolation (t(14) = 0.57, p = 0.58). **C)** For flies with a UAS enhancer sequence that induces RNAi targeted to *to* mRNA when activated, isolated females increase sucrose consumption relative to control females (t(21) = 3.41, p = 0.003). **D)** When RNAi targeted to *to* mRNA is induced in *to*-expressing cells, social isolation increases sucrose feeding in males (t(14) = 2.31, p = 0.04) yet decreases it in females (t(15) = 5.35, p < 0.0001) relative to respective controls. \**p* < 0.05, \*\**p* < 0.01, \*\*\*\**p* < 0.0001.

**S9 Fig. Overexpression of *to* in *to*-expressing cells results in no differences in sucrose feeding choice between isolated flies and their respective controls. A)** In ORE-R flies, our preliminary results indicated a female-specific increase in sucrose feeding following social isolation (t(15) = 3.14, p = 0.007). **B)** Flies expressing GAL4 under the control of a *to*-specific promoter do not show differences in sucrose feeding following social isolation. **C)** For flies with a UAS enhancer sequence that induces *to* expression when activated, isolated females increase sucrose consumption relative to control females (t(14) = 7.86, p < 0.0001). **D)** When *to* expression is induced in *to*-expressing cells, social isolation does not alter sucrose feeding in males (t(14) = 0.30, p = 0.77) or in females (t(15) = 1.59, p = 0.13) relative to respective controls. \*\**p* < 0.01, \*\*\*\**p* < 0.0001.

**S10 Fig. Knock down of *to* in neurons does not alter sucrose feeding across experimental groups. A)** In ORE-R flies, females increased their sucrose feeding following social isolation (t(15) = 3.14, p = 0.007). **B)** Flies expressing GAL4 under the control of an *elav*-specific promoter do not show differences in sucrose feeding following social isolation in either sex. **C)** For flies with a UAS enhancer sequence that induces RNAi targeted to *to* mRNA when activated, isolated females increase sucrose consumption relative to control females (t(21) = 3.41, p = 0.003). **D)** When RNAi targeted to *to* mRNA is induced in neurons, there is no difference in sucrose feeding choice between socially isolated and control flies of either sex. \*\**p* < 0.01.

**S11 Fig. Overexpression of *to* in neurons does not yield sucrose feeding differences between isolated and control flies of either sex. A)** In ORE-R flies, females increased their sucrose feeding following social isolation (t(15) = 3.14, p = 0.007). **B)** Flies expressing GAL4 under the control of an *elav*-specific promoter do not show differences in sucrose feeding following social isolation in either sex (males: t(16) = 0.30, p = 0.77; females: t(18) = 0.69, p = 0.50). **C)** For flies with a UAS enhancer sequence that induces *to* expression when activated, isolated females increase sucrose consumption relative to control females (t(14) = 7.86, p < 0.0001). **D)** When *to* expression is driven in neurons, there is no difference in sucrose feeding choice between socially isolated and control flies of either sex (t(9) = 0.87, p = 0.41). \*\**p* < 0.01, \*\*\*\**p* < 0.0001.

**S12 Fig. Isolated males and isolated females show no differences in sucrose-feeding relative to controls following knock down of *tra* in neurons. A)** In ORE-R flies, sucrose feeding is higher in isolated females compared to control females (t(15) = 3.14, p = 0.007). **B)** Flies expressing GAL4 under the control of an *elav*-specific promoter do not show differences in sucrose feeding following social isolation in either sex. **C)** For flies with a UAS enhancer sequence that induces RNAi targeted to *tra* mRNA when activated, isolated females (t(14) = 2.79, p = 0.01) and isolated males (t(14) = 3.91, p = 0.002) increase sucrose consumption relative to their respective controls. **D)** When RNAi targeted to *tra* is induced in neurons, there is no difference in sucrose feeding choice between socially isolated and control flies of either sex (males: t(17) = 1.42, p = 0.17; females: t(17) = 0.91, p = 0.38). \**p* < 0.05, \*\**p* < 0.01.

**S13 Fig. Downregulation and overexpression of *to* and downregulation of *tra* align with expectations across experimental groups. A)** We performed qPCR to confirm genetic knockdown of *to* in the to-Gal4 x UAS-to^RNAi^ flies and overexpression of *to* in to-Gal4 x UAS-to flies. *To* overexpression was clearer in control males and control females, which could be due to ORE-R isolated males and isolated females already having relatively high expression of *to*. **B)** Although the elav-Gal4 x UAS-to flies showed no change or negative expression of *to* relative to ORE-Rs, the elav-Gal4 x UAS-to^RNAi^ flies showed downregulation of *to* across experimental groups. **C)** Genetic knockdown of *tra* in elav-Gal4 x UAS-tra^RNAi^ flies decreased *tra* expression relative to ORE-Rs. **D)** Genetic knockdown of *tra* in elav-Gal4 x UAS-tra^RNAi^ flies also decreased expression of *to* across groups. **E)** In elav-Gal4 x UAS-to flies, *to* expression was elevated relative to elav-Gal4 flies in control males and isolated males but not in females. **F)** In elav-Gal4 x UAS-to flies, *to* expression was elevated relative to UAS-to flies across all experimental groups. After completing the VBFD test, flies were frozen on dry ice and decapitated. RNA was extracted from whole fly heads and reverse transcribed into cDNA. All groups represent n = 3 samples/group. Error bars are ± SEM.

**S1 Table. Average group size and number of groups tested in the VBFD test for all experimental groups across all genotypes.**

**S2 Table. List of PCR primers used for experiments.**

**S3 Table. Statistics for all 2x2 ANOVAs.**

**S4 Table. Statistics for all one-way ANOVAs.**

